# Ultrafast, one-step, and microwave heating-based synthesis of DNA/RNA-AuNP conjugates

**DOI:** 10.1101/2021.09.11.459879

**Authors:** Mengqi Huang, Erhu Xiong, Menglu Hu, Huahua Yue, Tian Tian, Debin Zhu, Xiaoming Zhou

**Author notes:** These authors contributed equally to this work (Mengqi Huang, Erhu Xiong).

## Abstract

DNA/RNA-gold nanoparticle (DNA/RNA-AuNP) nanoprobes have been widely employed for nanobiotechnology applications. Here we discovered that both thiolated and non-thiolated DNA/RNA can be efficiently attached to AuNPs to achieve high-stable spherical nucleic acid (SNA) within minutes under a domestic microwave (MW)-assisted heating-dry circumstance. Further studies showed that for non-thiolated DNA/RNA the conjugation is poly (T/U) tag dependent. Spectroscopy, test strip hybridization, and loading counting experiments indicate that low-affinity poly (T/U) tag mediates the formation of a standing-up conformation, which is distributed in the outer layer of such a SNA structure. In further applications study, CRISPR/Cas9-sgRNA (135 bp), RNA from Nucleocapsid (N) gene of SARS-CoV-2 (1279 bp), and rolling circle amplification (RCA) DNA products (over 1000 bp) could be successfully attached on AuNPs, which overcomes the routine methods in long-chain nucleic acid-AuNP conjugation, exhibiting great promise in novel biosensing and nucleic acids delivery strategy. This novel heating-dry strategy has improved the traditional DNA/RNA-AuNP conjugation methods in simplicity, rapidity, cost, and universality.

## Introduction

The invention of gold nanoparticles (AuNPs)-based spherical nucleic acids (SNAs), such as DNA-AuNP and RNA-AuNP conjugates, has attracted worldwide interest and opened up the field of nanobiotechnology^1-4^. This unique nanomaterial consists of AuNPs core and a dense nucleic acid, allowing molecular recognition and programmability of DNA/RNA to be combined with unique optical, chemical, electrical and catalytic properties of AuNPs, thus, impart SNA novel chem-physical properties and biological functions^5,6^. Numerous applications have since been made including programmed DNA/RNA nanotechnology, molecular diagnosis, imaging, gene regulation, and drug delivery^6-16^.

The construction of DNA-AuNP and RNA-AuNP conjugates is critical for these successful applications in biology, chemistry, medicine, and nanoscience. Conventionally, to attach negatively charged DNA and RNA to negatively charged AuNPs, a key is to screen the charge repulsion between AuNPs and nucleic acids. The most commonly used method is salt-aging method^17-19^, where salt was gradually added into the mixture of thiol (SH)- or poly (A)-tagged DNA and AuNPs to reduce charge repulsion. However, the salt-aging method required over 2 days to completion, making it an extremely time-consuming process. A few new labeling methods were developed by adding acids^20,21^, surfactants^22^, polymers^23,24^, and organic solvent^25^, however, the use of extra reagents complicates the labeling process and may reduce the applicability of SNAs. Recently developed freeze-thaw method could attach thiol-modified oligonucleotides on AuNPs in a time-effective and reagentless form^26^. However, DNA/RNA thiolation increases the cost, especially for long oligonucleotides, whose chemical modification is extremely expensive and difficult ^27,28^. Our group have successfully used a freeze-thaw method to attach thiol-free DNA/RNA on AuNPs, but this method still lacks universality due to sequence structure effect^29^. In summary, it seems that there is still a lack of an ideal DNA-AuNP and RNA-AuNP construction method that can simultaneously satisfy the simplicity, rapidity, low cost, and versatility. In particular, almost all currently reported DNA-AuNP and RNA-AuNP construction methods only demonstrated their feasibility for functionalization of short DNA and RNA sequences. Long DNA/RNA-conjugated AuNPs nanostructures, which may be developed as potential new biosensing, DNA/RNA vaccine^30,31^, and gene editing tool delivery strategies^32,33^, have been rarely developed.

Herein, we reported a microwave (MW)-assisted heating-dry method to construct SNAs, where a domestic microwave oven was used as a heater to drive the labeling process to completion in minutes without the need for extra reagents, and demonstrated its applicability to both thiolated and non-thiolated DNA/RNA. Further, for attaching non-thiolated DNA/RNA, we found that DNA/RNA-AuNP conjugates are poly (T/U) tag dependent under heating-dry circumstance. Spectroscopy, test strip hybridization, and loading counting experiments indicate that poly (T/U) tag is distributed in the outer layer of such a SNA structure. It is inferred that this essential low-affinity poly (T/U) tag is mainly responsible for the formation of a “standing-up” conformation in the AuNPs surface. Notably, our method addresses the deficiencies of traditional methods in long DNA/RNA labeling and allows selective functionalization of AuNPs with sgRNA (135 bp), RNA (1279 bp) from Nucleocapsid (N) gene of SARS-CoV-2, and rolling circle amplification (RCA) DNA products (over 1000 bp). In addition, the potential applications of this labeling strategy were demonstrated by developing new colorimetric identification of DNA single-base mutation or detecting viral DNA based on CRISPR-based lateral flow strategy. The successful construction of long and structured DNA/RNA-conjugated AuNPs nanostructure may also indicate that a new functional nucleic acids delivery system based on AuNPs as a carrier can be developed. Overall, currently presented labeling strategy represents an ultrafast, simple, cost-effective and universal method that can be extended to the conjugation of almost all types of DNA and RNA sequences, which will expand the use of SNAs in biomedicine and nanotechnology in the future.

## Results and discussion

### Ultrafast construction of DNA-AuNP and RNA-AuNP conjugates based on MW-assisted heating-dry method

Both nucleic acids and AuNPs solutions exhibit negatively charged properties, in order to promote effective nucleic acid-AuNP conjugation, some well-proven strategies, such as salt-aging^17,18^ and low-pH methods^20,21^, are to shield the charge to benefit the affinity interaction between nucleic acids and AuNPs. Other effective strategies, such as freeze-thaw^26^ and solvent methods^25^, are to increase the local concentration by compressing the physical space, thereby forcing the chemical covalent reaction or physical cross-linking between AuNPs and nucleic acids in an ultra-localized reaction volume. Our previous study has shown that freeze-thaw method encounters the influence of secondary structure of oligonucleotides on AuNPs-based labeling reaction^29^, thus lacks sequence universality for construction of AuNPs-based bioprobes. Here, we invented a new DNA-AuNP and RNA-AuNP construction strategy through MW-assisted heating-dry method (Fig. 1a, b). Heating is well known to destroy the higher-order nucleic acid structure and stretch the nucleic acid strands^34^. The microwave oven, which is used in almost every kitchen, is a simple and cost-effective heating device^35,36^. In particular, the microwave radiation can be completely reflected on the metal surface and therefore has no effect on the metal particles, while the aqueous solution can be quickly heated and dried. We envisioned that this heating-dry strategy could be an ideal DNA/RNA-AuNP labeling method due to its role in simultaneously stretch the nucleic acid structure and concentrate the reaction volume. Accordingly, we firstly mixed SH-DNA with AuNPs solution (Supplementary Fig. 1). Surprisingly, after a few minutes of MW heating and drying, AuNPs showed red clumps, while AuNPs showed purple-black clumps with thiol-free DNA or in the absence of DNA (Supplementary Fig. 1). After addition of salt solution to the resulted red clumps, AuNPs immediately redispersed into a red colloidal state, indicating that AuNPs and SH-DNA may be covalently conjugated (Supplementary Fig. 1). These phenomena signified that a simple DNA-AuNP labeling technique using microwave-assisted heating-dry method has been developed.

**Fig. 1.**
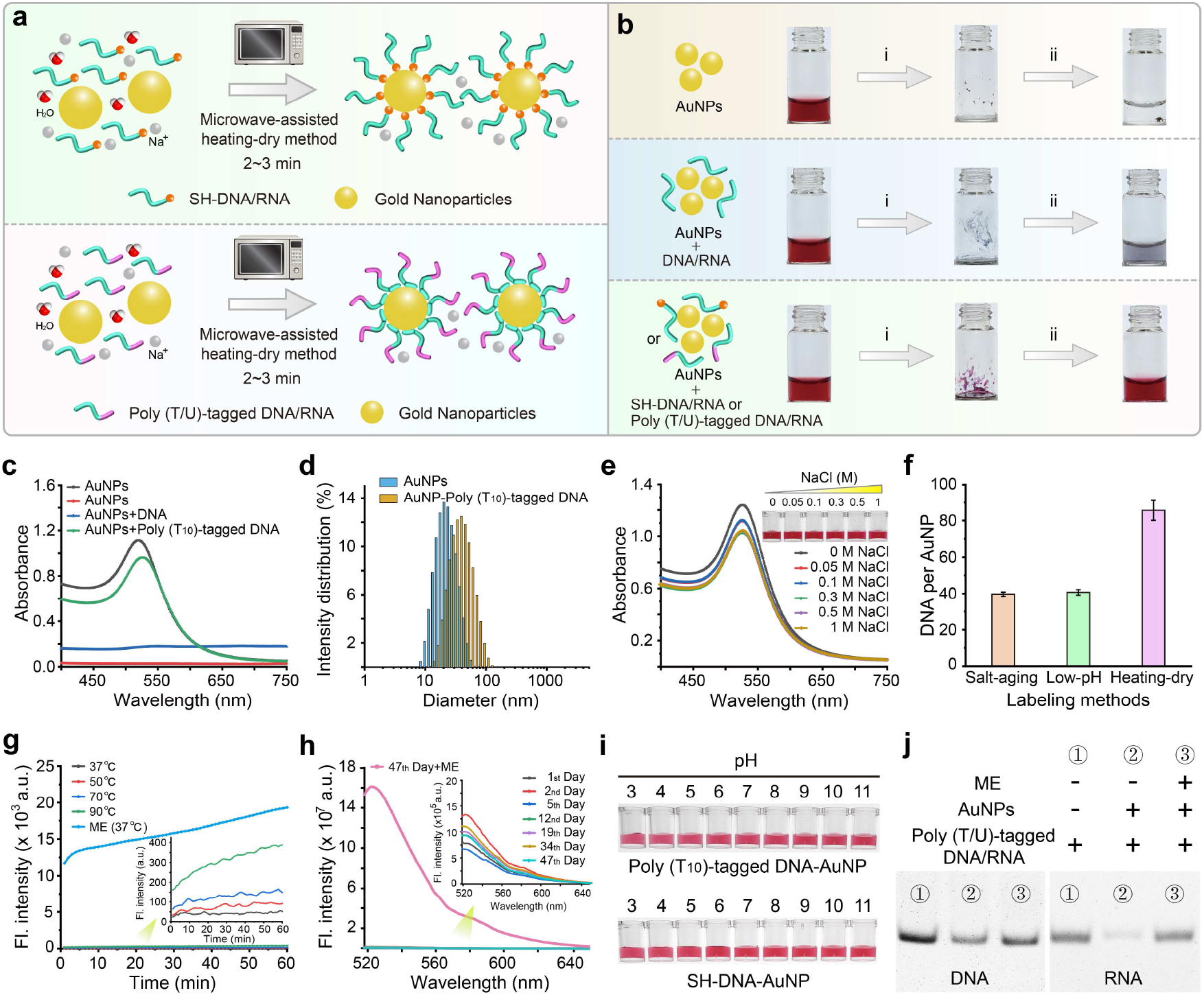
Ultrafast construction of DNA/RNA-AuNP conjugates based on MW-assisted heating-dry method. **(a)** Scheme of attaching thiolated and non-thiolated DNA/RNA to AuNPs using MW-assisted heating-dry method, in which the heating-dry process is driven by a domestic microwave oven for 2∼3 min. **(b)** Photographs of AuNPs before and after heating-dry process. The bare AuNPs and random DNA/RNA sequence mixed AuNPs aggregated after heating dry treatment. While poly (T/U)-tagged DNA-AuNP retained monodispersed and seems red after heating dry and resuspension. i: MW-assisted heating dry; ii: resuspending with water or buffer. **(c)** Characterization of the non-thiolated DNA/RNA-AuNPs conjugates based on MW-assisted heating-dry labeling method using UV-vis absorption spectroscopy. **(d)** Dynamic light scattering measurement of the hydrodynamic diameters of bare AuNPs and DNA-AuNP conjugates. **(e)** Photographs and absorption spectroscopy showing the stability evaluation of DNA-AuNP conjugates in different NaCl concentrations. **(f)** The measured number of DNA probes attached to each AuNP. The DNA-AuNP conjugates were constructed using poly (A)-based salt-aging and low-pH methods, and poly (T)-based MW-assisted heating-dry method, respectively. **(g)** Thermal stability evaluation of DNA-AuNP conjugates. The process of DNA desorption from AuNPs was monitored by measuring the fluorescence intensity of FAM on DNA probes at varied temperatures. **(h)** Time stability evaluation of DNA-AuNP conjugates. The process of DNA dissociation from AuNPs was monitored by measuring the fluorescence intensity of FAM on DNA probes at different storage times. **(i)** pH stability evaluation of thiolated and non-thiolated DNA-AuNP conjugates. **(j)** Evaluation of DNA/RNA integrality of the resulted DNA/RNA-AuNP conjugates.

We further seek to develop a universal and thiol-free nucleic acid-AuNP construction method (Fig. 1a). Although thiol modification is very easy to obtain for short nucleic acid sequences, for long-chain DNA and RNA, the chemical modification is very expensive and difficult. Therefore, conjugation of long-chain DNA and RNA to AuNPs is rarely achieved. Adenine (A) base has been well confirmed to have good affinity for Au^37,38^, and poly (A)-tagged nucleic acid sequences have also been successfully applied to the coupling of AuNPs based on freeze-thaw or salt-aging methods^18,29^. Therefore, we take it for granted that poly (A)-tagged nucleic acid sequences should be suitable for the heating-dry labeling strategy we developed. However, to our surprise, in our attempt we found that only the poly (T/U)-tagged nucleic acids showed a preferred labeling efficiency under heating-dry circumstance (Fig. 1a, b). In the absence of DNA/RNA sequences or in the presence of random DNA/RNA sequences without poly (T/U) tags, AuNPs solutions will aggregate irreversibly (Fig. 1b). Further, results showed that heating-dry strategy can also be applied to labeling of larger-sized AuNPs and SH-DNA-gold nanorods (AuNRs) conjugation (Supplementary Fig. 2).

In order to further characterize this DNA-AuNP conjugate, we directly measured the UV-vis spectroscopy of the poly (T)-tagged DNA conjugated AuNPs solution. It was shown that the absorption peak of AuNPs with poly (T)-tagged DNA exhibited a 6 nm red-shift, suggesting that there are DNA attached on the AuNPs surface (Fig. 1c). To further demonstrate the poly (T)-tagged DNA attachment on AuNPs, the AuNPs solution was centrifuged and washed and then analyzed by dynamic light scattering (DLS). The hydrated particle size of DNA-AuNP conjugate is larger than the bare AuNPs (Fig. 1d and Supplementary Table 1), further indicating that poly (T)-tagged DNA strands were attached. In addition, the electron microscope results confirmed that the MW-assisted heating treatment did not damage the morphology of AuNPs (Supplementary Fig. 3).

Next, we wonder whether the DNA-AuNP conjugate is enough stable. Firstly, we tested its stability by resuspending it with different salt concentrations. Results showed that the DNA-AuNP conjugates were still monodispersed even challenge 1 M NaCl solution and its absorption peak did not show observable shift (Fig. 1e), while the bare AuNPs aggregated at a salt concentration of 50 mM (Supplementary Fig. 4). Such high salt stability indicated that there were many poly (T)-tagged DNA sequences attached to the AuNPs surface. We subsequently measured the number of FAM-labeled poly (T)-DNA probes assembled on each AuNP according to a β-mercaptoethanol (ME)-based displacement experiment (Supplementary Fig. 5a-c)^39^. As a comparison, the number of FAM-poly (A)-DNA on each AuNP that labeled using two reported thiol-free methods, poly (A)-based salt-aging and low-pH methods, were also measured. Results showed that the number of poly (T)-tagged DNA probes attach to each AuNP by MW-assisted heating-dry method is 83, which is about 2 times higher than the salt-aging and low-pH method (Fig. 1f). Further test indicated that when SH-DNA was used, for each AuNP approximately 270 DNA strands can be loaded (Supplementary Fig. 5d). We believe that the high DNA density is not due to the multilayered adsorption that caused by hydrogen bonding between DNA strands. This is confirmed in a subsequent thermal stability experiment, where FAM-labeled DNA-AuNP conjugates exhibited negligible fluorescence increase in the solution at different temperatures, indicating high thermal stability of the DNA-AuNP conjugates and non-specific adsorption was also excluded (Fig. 1g). However, a rapid increase in fluorescence intensity was observed when the replacement reagent ME was mixed with the DNA-AuNP conjugates (Fig. 1g). At the same time, we also monitored the time stability based on the measurement of the fluorescence of DNA-AuNP conjugates. The results showed that only negligible fluorescence intensity increases even for up to 47-day storage, indicating that the conjugates are stable for long-time storage (Fig. 1h). Further, the DNA-AuNP conjugates stability at a wide pH range (pH 3∼11) was also tested. It was showed that both thiolated and non-thiolated DNA/RNA-AuNP conjugates exhibit excellent pH tolerance (Fig. 1i).

One may worry about whether such a high-temperature labeling condition will damage DNA and affect subsequent use. We then tested the nucleic acid integrity and hybridization ability of the resulted DNA/RNA-AuNP conjugates. It was found that DNA/RNA are still intact and the conjugates work as well as the conventional SH-DNA-labeled AuNP probes based on a test strip hybridization experiment (Fig. 1j and Supplementary Fig. 6).

### Non-thiolated DNA/RNA-AuNP conjugation is poly (T/U) tag dependent

To further understand the sequence-based labeling effect, we designed four types of DNA with poly (A_10_), poly (T_10_), poly (C_10_), and poly (G_10_) tags at 5′- and 3′-terminal and mixed them with AuNPs, respectively. Moreover, four RNAs with poly (rA_10_), poly (rU_10_), poly (rC_10_), and poly (rG_10_) tag at 3′-terminal were also applied. After heating-dry and centrifugation-washing treatment, the DNA/RNA-AuNP conjugates were resuspended in 1 M NaCl solution. Results showed that only AuNPs with poly (T/U)-tagged DNA/RNA retained red color in 1 M NaCl solution, indicating AuNPs were protected by high-density DNA strands (Fig. 2a, i, ii, iv). Further assay showed that poly (A/T/C/G) tags at the middle position exhibited the similar phenomena (Fig. 2a, iii). Labeling using DNA sequences with poly (A/T/C/G) tags with varied length have also been tested. Under all situations only these poly (T)-tagged DNAs are efficient, thus the poly (A/C/G) tags length effect is excluded (Fig. 2b and Supplementary Fig. 7). The results also showed that five T bases appeared to be enough for efficient labeling when the T bases are located at 5′- and 3′-terminal, and more than 10 T bases did not significantly improve the labeling efficiency (Fig. 2b). For T base located at the middle region of the DNA sequence, the labeling efficiency increased with the increase of T base number and T_30_ base showed the highest labeling efficiency (Fig. 2b). All these observations indicated that poly (T) tag is essential for successful labeling and this labeling method exhibits flexibility in the position and length of poly (T) tag. In addition to the DNA sequence with terminal and middle tags, we also proved that the MW-assisted heating-dry method can achieve successful labeling for DNA sequence with poly (T) tag located at both ends or hairpin DNA sequence with shielded poly (T) tag in the stem region (Supplementary Fig. 8a, b). The results indicated that heating-dry method can be applied to structured nucleic acid labeling, thus providing a more generic way than previously reported techniques, such as freeze-thaw method (Supplementary Fig. 8c).

**Fig. 2.**
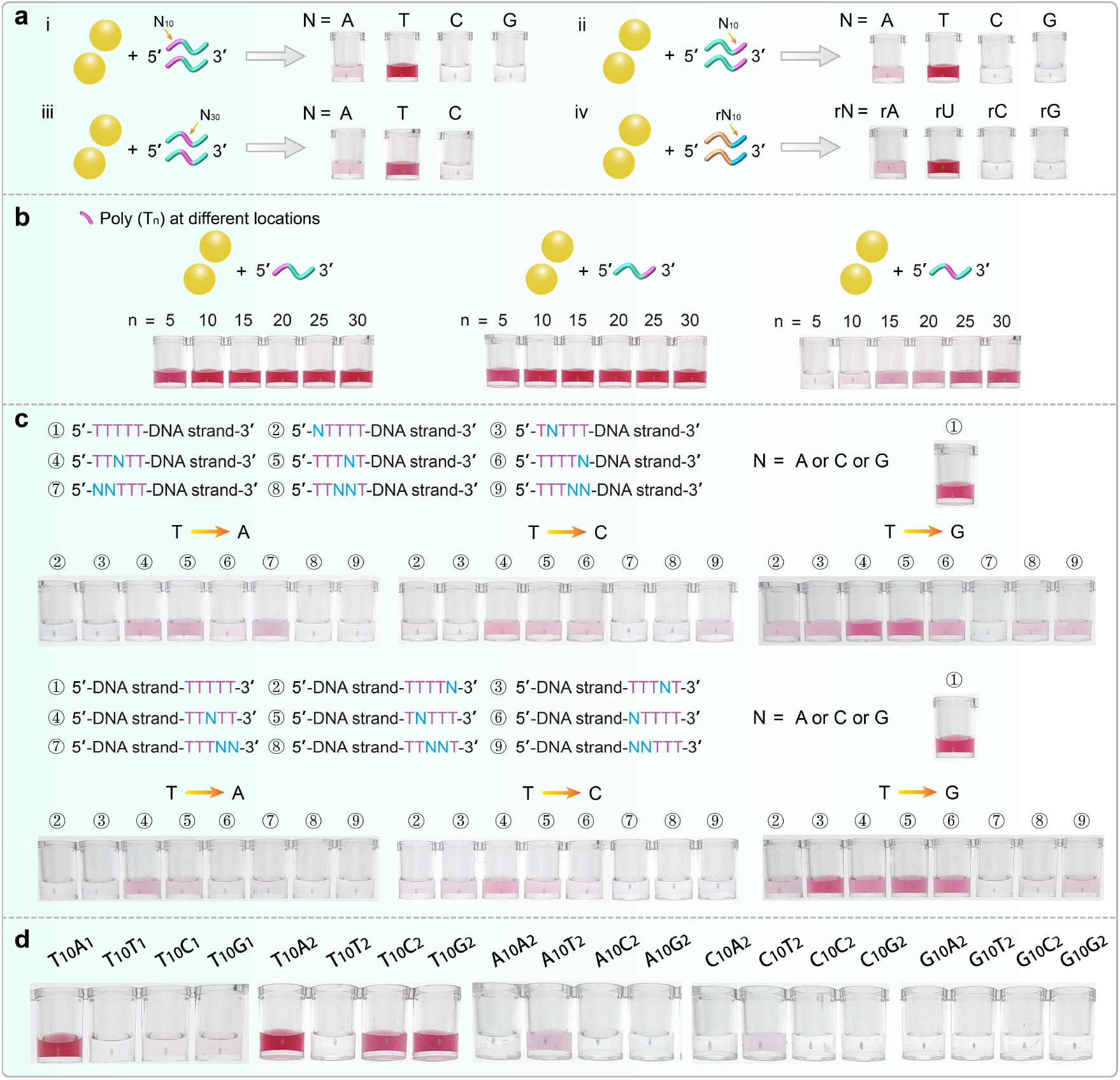
Poly (T/U) tag-dependent DNA/RNA-AuNP conjugates. **(a)** MW-assisted heating-dry labeling of DNA-AuNP and RNA-AuNP conjugates. Photographs showing the labeling results of DNA sequences with poly (T) tag at 5′-terminal (i), 3′-terminal (ii), and in the middle region (iii). Photographs showing the labeling results of RNA sequence with poly (rU) tag at 5′-terminal (iv). An inefficient and failed labeling will lead to the AuNPs aggregation. **(b)** Photographs showing the labeling results of poly (T) tagged-DNA with different numbers of T base located at 5′-terminal, 3′-terminal and in the middle region. **(c)** Photographs showing the labeling results of poly (T_5_)-tagged DNA, in which one or two T bases in T_5_ were replaced by A, C, and G base. Poly (T) tags at both 5′- and 3′-terminal are evaluated. **(d)** Observation of the poly (T_10_) dependence and role in the heating-dry labeling process.

To more detailly evaluate the role of T base in labeling process, we replaced one or two T bases with A, C, or G base in poly (T_5_) -tagged DNA sequences. Results showed that in most case the labeling efficiency dramatically decreased or totally lost upon this replacement (Fig. 2c), which indicated that five consecutive T bases are very critical. We also noticed that when replacing single T with G the labeling efficiency did not dramatically decreased. Previous observations and our colorimetric experiments indicated that T base has the lowest Au affinity (Supplementary Fig. 9). Some pioneering works have concluded that the Au affinity order for all four bases is: A > C > G > T^40-42^. Dependence of low-affinity T base during the heating-dry process prompted us to know how it is involved in this labeling method. To simplify the base effect evaluation, we designed four types of sequences consisting of two kinds of bases to observe this labeling response. Interestingly, we found that the T_10_A_1_ sequence was effectively conjugated, while T_10_T_1_, T_10_C_1_, and T_10_G_1_ did not (Fig. 2d). Further experiments showed that T_10_A_2_, T_10_C_2_, and T_10_G_2_ can also be effectively conjugated while T_10_T_2_ does not (Fig. 2d). However, when A_10_, C_10_ and G_10_ were used as the tag, all labeling reactions were unsuccessful (Fig. 2d). These experiments implied that T base is essential in the heating-dry labeling strategy but it is not directly involved in the DNA-AuNP conjugation.

### Experimental observation of poly (T) tag orientation

In order to clarify the underlying labeling mechanism, we firstly used surface enhanced Raman spectroscopy (SERS) to observe the poly (T) tag orientation in the SNA structure. Because the labeling mechanism based on Au-SH conjugates is well proved, we then tested the SERS signal based on the hybridization of a 5′-SH-DNA-AuNP conjugates and a complementary DNA (cDNA) probe with ROX labeled at 5′- and 3′-terminal, respectively. When 3′-ROX-cDNA-5′ and 3′-cDNA-ROX-5′ hybridized with the 5′-SH-DNA-AuNP conjugates, respectively, ROX will be close to or away from AuNPs surface. Expectantly, we observed significant SERS signal enhancement after 3′-ROX-cDNA-5′ probe hybridization (Fig. 3a, i). This indicated that SERS is suitable for observing the poly (T) tag orientation. Accordingly, we designed an “amphiphilic” DNA sequence with A_5_ and T_5_ tags located at 5′- and 3′- terminals, respectively, for AuNP conjugation. 3′-ROX-cDNA-5′ and 3′-cDNA-ROX-5′ probes were used for observing the SERS signals after hybridization. Experiments showed that 3′-ROX- cDNA-5′ probe mediated significant SERS signal enhancement (Fig. 3a, ii). When replaced the T_5_ tag with T_10_ tag, more significant SERS signal was observed (Fig. 3a, iii). These results supported that poly (T) tag is distributed in the outer layers of SNA structures.

**Fig. 3.**
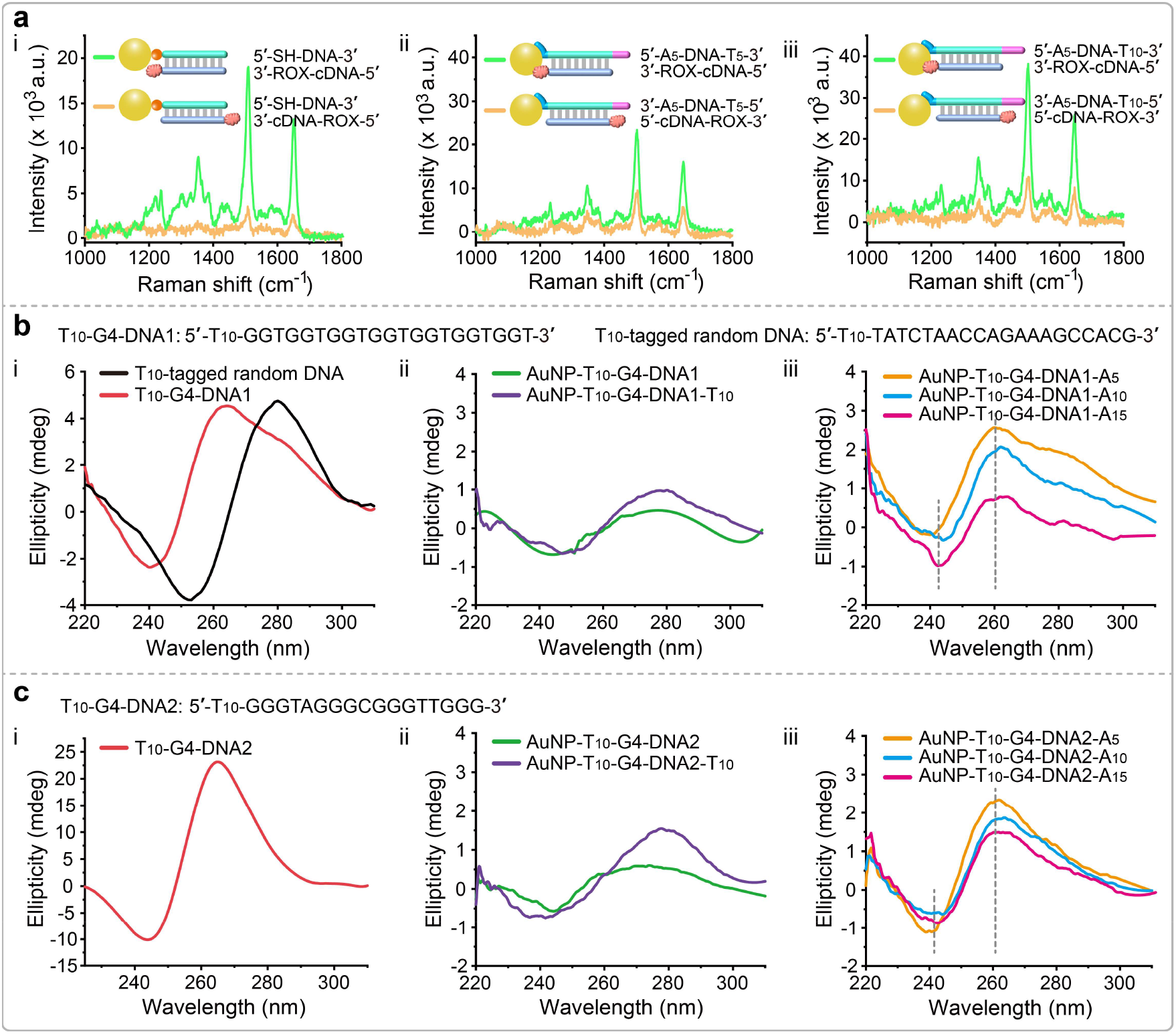
Spectroscopic observation of poly (T) tag orientation in the SNA structure. **(a)** i: SERS experiments showing the Raman signal variation when cDNAs labeled with ROX at 5′- and 3′-terminal hybridized with the SH-DNA-AuNP conjugates, respectively. ii: SERS experiments showing the Raman signals variation when cDNA labeled with ROX at 3′-terminal hybridized with the A_5_-DNA-T_5_ and T_5_-DNA-A_5_ conjugated AuNP probes, respectively. iii: SERS experiments showing the Raman signal variation when cDNA labeled with ROX at 3′-terminal hybridized with the A_5_-DNA-T_10_ and T_10_-DNA-A_5_ conjugated AuNP probes, respectively. For all SERS experiments, 30 nm diameter AuNPs are used. **(b)** i: CD spectra of the T_10_-tagged G4-DNA1 sequence and T_10_-tagged random DNA sequence. G4-DNA1 sequence has two characteristic peaks at around 240 and 260 nm. Random DNA sequence has two characteristic peaks at around 250 and 280 nm. ii: CD spectra of AuNP-T_10_-G4-DNA1 and AuNP- T_10_-G4-DNA1-T_10_ conjugates. iii: CD spectra of AuNP-T_10_-G4-DNA1-A_5_, AuNP-T_10_-G4-DNA1-A_10_, and AuNP-T_10_-G4-DNA1-A_15_ conjugates. **(c)** i: CD spectra T_10_-tagged G4-DNA2 sequence. ii: CD spectra of AuNP-T_10_-G4-DNA2 and AuNP-T_10_-G4-DNA2-T_10_ conjugates. iii: CD spectra of AuNP-T_10_-G4-DNA2-A_5_, AuNP-T_10_-G4-DNA2-A_10_, and AuNP-T_10_-G4-DNA2-A_15_ conjugates.

We further executed a structured DNA labeling experiment to elucidate labeling mechanism. Two kinds of DNA sequences with well-confirmed G-quadruplex (G4) structures (G4-DNA1 and G4-DNA2) are employed. The circular dichroism (CD) spectroscopy was used to measure these two G4-contained sequences and random sequence, and the results showed G4-DNA1 and G4-DNA2 contained sequences present characteristic peaks at around 240 and 260 nm while random DNA sequence presents characteristic peaks at around 250 and 280 nm (Fig. 3b, i and Fig. 3c, i). Subsequently, we designed G4-DNA1 with T_10_ tag at 5′-terminal and G4-DNA2 with T_10_ tag at both 5′- and 3′-terminal for the AuNP conjugation. According to this design, it can be expected that if T_10_ is involved in the AuNP conjugation, the G4 sequence will expose to the outer layer so that its structure is retained. Conversely, if the G4 sequence is involved in the AuNP conjugation, the G4 structure will be destroyed. CD spectra showed that G4 characteristic peaks disappeared after AuNP conjugation, verifying that G4 sequence is involved in the AuNP conjugation (Fig. 3b, ii and Fig. 3c, ii). Further, we replaced the T_10_ tag at 3′-terminal with A_5_, A_10_, and A_15_ tags, respectively. It is expected that high-affinity poly (A) tag will preferentially bind to the AuNPs surface, so the G4 sequence will be released to maintain its intact structure. CD spectra of these resulted DNA-AuNP conjugates are tested, which showed that G4 characteristic peaks are restored (Fig. 3b, iii and Fig. 3c, iii).

In order to strengthen the understanding of labeling mechanism, and also supply guide for future applications, we employed test strip hybridization assay to evaluate the hybridization properties of the resulted DNA-AuNP conjugates. DNA sequences with varied poly (T) tag lengths were designed for AuNP conjugation and a cDNA can hybridize with these DNA sequences was pre-coated on the test strip. It was found that with the increase of poly (T) tag length, the hybridization efficiency decreased, whether 13 nm or 30 nm AuNP bioprobes (Fig. 4a, i). These results indicated that increasing the poly (T) length leads to an increase in steric hindrance and a decrease in hybridization efficiency. Subsequently, the same DNA sequences were tagged with a fixed poly (T_7_) at 5′-terminal and varied poly (A) lengths at 3′-terminal. The resulted DNA-AuNP conjugates were applied to the test strip hybridization and presented almost equal hybridization efficiency when increasing the poly (A) length from A_2_ to A_15_ (Fig. 4a, ii). It was also noticed that further increase of poly (A) length to 25 has a reverse effect on hybridization (Fig. 4a, ii). It is possible that the longer poly (A) tag leads to a decrease in the number of probes conjugated to the AuNPs surface, thus affecting the hybridization. At last, we pre-coated a poly (A_7_) probe on the test strip to test the above-mentioned DNA-AuNP conjugates. Robust and equal hybridization efficiency can be observed for DNA-AuNP conjugates with varied poly (A) lengths (Fig. 4a, ii). These experiments provide strong evidence that poly is not involved in conjugation with AuNPs and is distributed in the outer layer of the SNA structure. This can also be further corroborated by measuring the attached DNA numbers at three kinds of labeling situations, where Poly (T_n_)-Poly (C_5_)-FAM, Poly (T_10_)-Poly (C_n_)-FAM, and Poly (T_10_)-Poly (C_5_)-FAM were used for labeling (Supplementary Figs. 10-12). Attached DNA is almost unchanged with varied low-affinity poly (T) lengths but maintains fixed high-affinity poly (C_5_) tag (Fig. 4b, i). Keep the poly (T_10_) tag fixed, while increasing the length of poly (C) and poly (A) tags, attached DNA is gradually reduced (Fig. 4b, ii and iii).

**Fig. 4.**
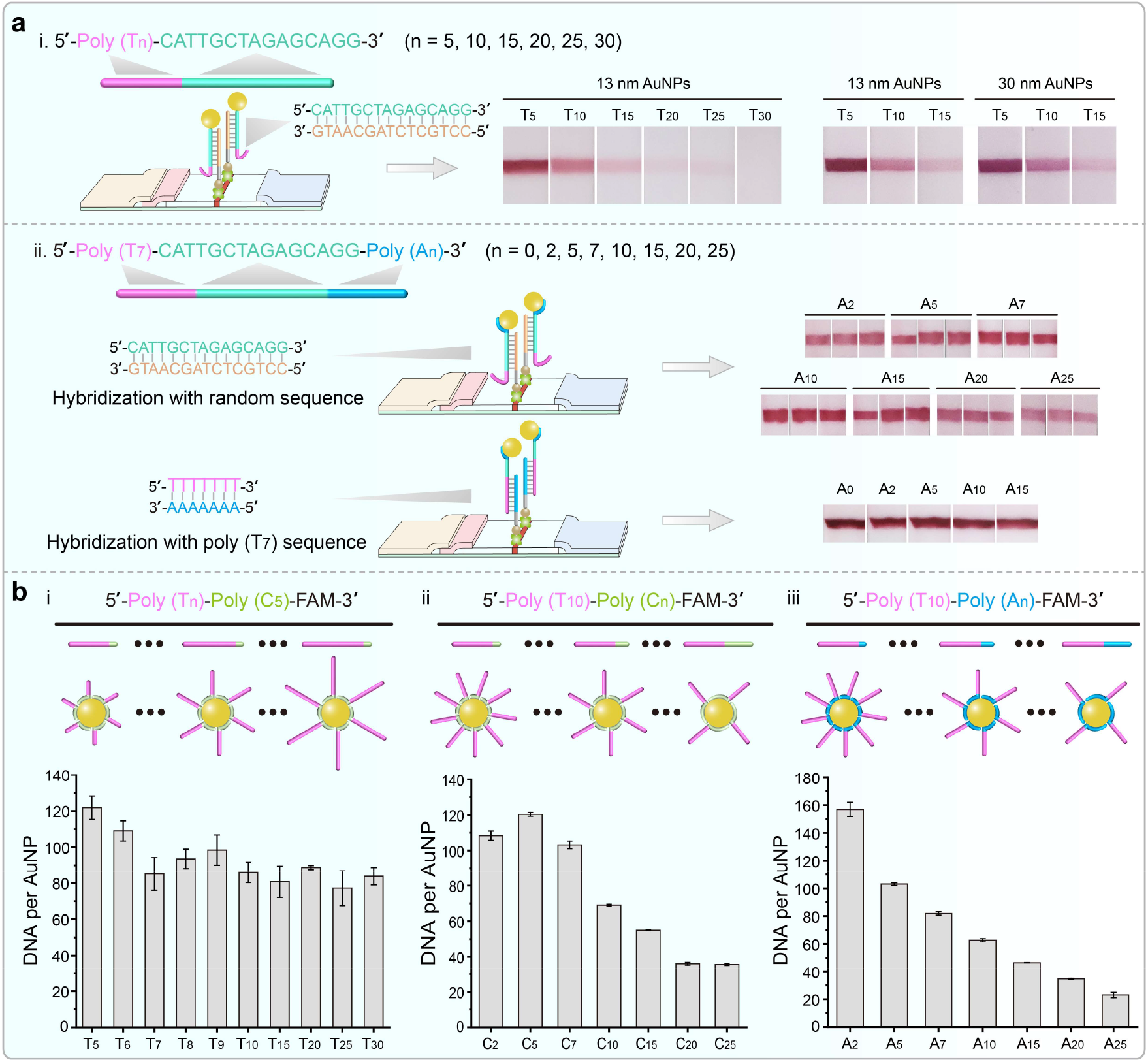
Verification of poly (T) tag orientation in the SNA structure using test trip hybridization and loading counting experiments. **(a)** i: Scheme showing the poly (T_n_)-tagged DNA probes are used for AuNPs labeling and the lateral flow hybridization device. A complementary DNA probe is pre-coated on the lateral flow device used for hybridization. 13 and 30 nm diameter AuNPs are used for labeling and subsequent hybridization experiments. ii: DNA sequences with poly (T_7_) and poly (A_n_) tags at both terminals are used for AuNPs labeling and subsequent hybridization experiments. Two kinds of DNA probes are pre-coated on the lateral flow device for hybridizing with T_7_ and random DNA region, respectively. Hybridization results are observed by the naked eyes. **(b)** Counting of the attached DNA number from three labeling situations. Three kinds of DNA probes including (i) Poly (T_n_)-Poly (C_5_), (ii) Poly (T_10_)-Poly (C_n_), and (iii) Poly (T_10_)-Poly (C_5_), are used for labeling. FAM dye is modified at 3′-terminal of these DNA sequences.

### Mechanism description

It is fascinating that poly (T/U)-tagged DNA/RNA can be loaded on the AuNPs surface at high density by simply heating-dry treatment. Previous study has shown that heating could drive DNA to be adsorbed onto AuNPs or graphene^43,44^. To understand whether this labeling is only induced by heating, some random DNA sequences with poly (A_10_/T_10_/C_10_/G_10_) tags at 5′-terminal were mixed with AuNPs under heating (Fig. 5a and Supplementary Fig. 13). As observed, under both 90 and 150 °C, the AuNPs aggregated after the centrifugation-resuspension treatment, which show that heating process cannot mediate efficient DNA-AuNP conjugation (Fig. 5a, top panel and Supplementary Fig. 13). Conversely, all poly (T)-tagged DNA sequences could be efficiently attached to AuNPs using heating-dry method (Fig. 5a, below panel). In another test, when AuNPs solution were mixed with six kinds of DNAs and dried at lower temperatures (30 and 90 °C), feasible labeling was achieved although with uniform labeling efficiency (Supplementary Fig. 13). The uniform labeling efficiency may be due to the structural influence of some DNA sequences. Conversely, high and uniform labeling efficiency can be achieved only when both drying and heating conditions are available (Supplementary Fig. 13).

**Fig. 5.**
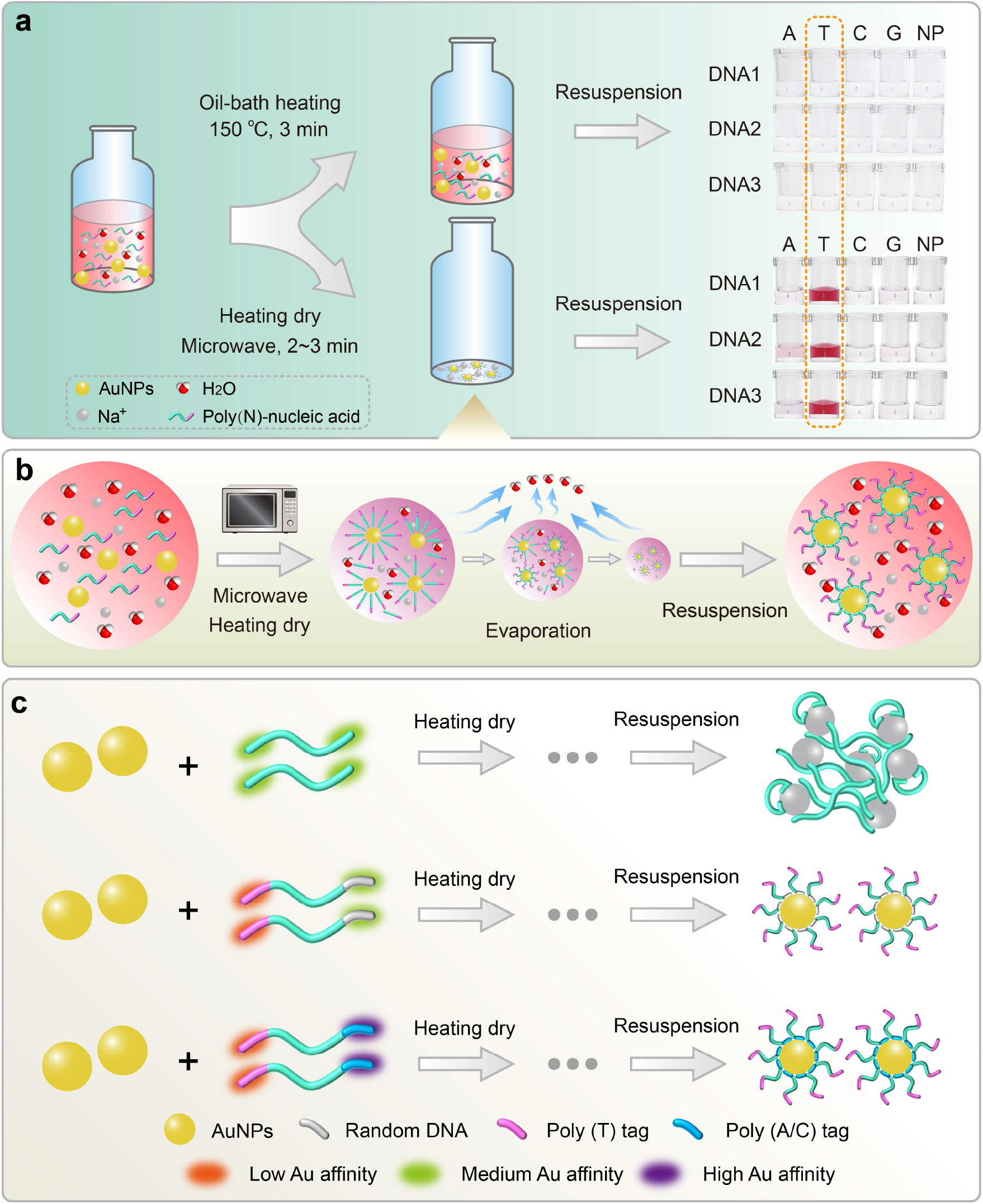
Mechanism description on DNA-AuNP conjugation by the MW-assisted heating-dry method. **(a)** Photographs showing a failed labeling with solely heating at 150 °C for 3 min, while there was a successful labeling of poly (T)-tagged DNA with heating-dry method. Poly (A/C/G)-tagged DNA and non-poly base (NP) DNA probes exhibited failed labeling. An inefficient and failed labeling will lead to the AuNPs aggregation. **(b)** MW-assisted heating-dry method involves two consecutive processes: i, MW-assisted heating induced high temperature unwinds the coiled structure of oligonucleotides. ii, MW-assisted water evaporation concentrates the DNA/RNA, AuNPs and salt, thereby accelerating the attachment of DNA/RNA on the AuNPs surface. **(c)** DNA sequences without poly (T) tag present medium Au affinity at both terminals. Under heating-dry conditions, DNA adsorbed on the AuNPs surface disorderly. Inadequate DNA adsorption or DNA-mediated crosslink adsorption will cause the AuNPs aggregation. Benefiting from the low Au affinity of poly (T) tag, poly (T)-tagged DNA exhibits an “amphiphilic”-like characteristic. Therefore, the sequence at one end (either a random sequence with medium Au affinity or poly (A/C) sequence with high Au affinity) mediates the DNA-AuNP conjugation, and the poly (T)-contained sequence at the other end is distributed in the outer layer of SNA structure.

Further, we found that poly (T_10_)-tagged DNA at the concentration of 1.25 µM is enough for heating-dry labeling (Supplementary Fig. 14a). Based on the data of DNA loading per AuNP (83 poly (T_10_)-tagged DNA strands each AuNP) and the measured AuNPs concentration after labeling completion, we were surprised that more than 80% of the DNA strands were utilized in the labeling process. Such a high DNA utilization ratio should be ascribed to the concentration effect under the heating-dry condition. For SH-DNA, a critical DNA concentration of 5 µM is required for successful conjugation (Supplementary Fig. 14b), which is consistent with the results of about 270 SH-DNA strands attached for each AuNP (Supplementary Fig. 3d). These observations indicated that both evaporation-mediated DNA condensing and heating-mediated DNA stretch cooperatively promote the high-density DNA loading on AuNPs surface (Fig. 5b).

Based on the above observations and poly (T) tag orientation experiments, we concluded that the heating-dry method presents the following mechanism: i. for random DNA/RNA sequence, both 5′- and 3′-terminal or middle region have medium and undifferentiated Au affinity. Under rapid heating-dry condition, flat and cross-linked DNA-AuNP adsorption behavior is dominant. The resulted DNA-AuNP complex is thus unstable against heating and high salt concentration due to the low DNA adsorption density (Fig. 5c); ii. poly (T/U)-tagged DNA/RNA exhibits an “amphiphilic”-like characteristic, where poly (T/U) tag has low Au affinity and the other end has medium Au affinity (random base composition) or high Au affinity (A/C-rich DNA sequences). Under rapid heating-dry condition, nucleic acid sequences are clustered and medium- and high-affinity regions are preferentially bound to AuNPs, thus the low-affinity poly (T/U)-contained region is distributed in the outer layer due to the repulsion effect. Such a “standing-up” conformation allows high-density nucleic acid strands attached thus mediating the formation of high-stable SNA structure (Fig. 5c). It is important to point out that for efficient conjugation the low-affinity tag is essential from all our observations. In a test, labeling failure occurred when the Au affinity of poly (T) was enhanced by phosphorothioate (PS) modification. However, addition of modification-free poly (T) tag at the other end, the successful conjugation was restored (Supplementary Fig. 15).

### MW-assisted heating-dry method enables colorimetric detection of poly (T) interspaced RCA products

After establishing an ultrafast, low-cost and facile DNA-AuNP construction method, we wondered whether this method could be used for long-stranded DNA-AuNP conjugation. RCA, an isothermal amplification reaction, is performed using circular DNA template and DNA primer to produce long single-stranded DNA (ssDNA) which contains many tandem repeating sequences complementary to circular DNA template^45^. Here, we designed a padlock probe containing a A_40_ sequence at non-recognition region and a 20 bp sequence for specific target DNA recognition. After performing ligation and RCA reaction, the resulting long ssDNA products will contain interspaced tandem repeated T_40_ sequence, which is potential suitable for AuNP-based labeling using heating-dry method (Fig. 6a). The gel analysis showed that the length of RCA products is over 1000 bp (Fig. 6b). With MW-assisted heating-dry method, we observed that the poly (T)-interspaced RCA products could be effectively attached thus AuNPs presented in a red monodispersed solution, while AuNPs aggregated when mixed with RCA products with random repeated sequence (Fig. 6c), which indicated a selective AuNPs-based attachment with T_40_ sequence in RCA products. Based on this observation, we further developed a simple colorimetric single-base mutation diagnostic platform. As shown in Fig. 6d, the padlock probe will be ligated in the present of perfectly matched target sequence thus the RCA reaction could generate long ssDNA strands containing T_40_ sequence, while no ligation and RCA amplification occur with single-base mismatched sequence (Fig. 6e). Subsequently, heating-dry method was utilized to colorimetric detection of the RCA products. The solution retained red when the sample with perfectly matched target, while it was colorless when the sample with single-base mismatched sequence or without target due to AuNPs aggregation (Fig. 6f). These results indicated that our developed diagnostic platform based on padlock-dependent ligation, RCA reaction and MW-assisted heating-dry method enables a new colorimetric gene mutation detection of DNA sequence.

**Fig. 6.**
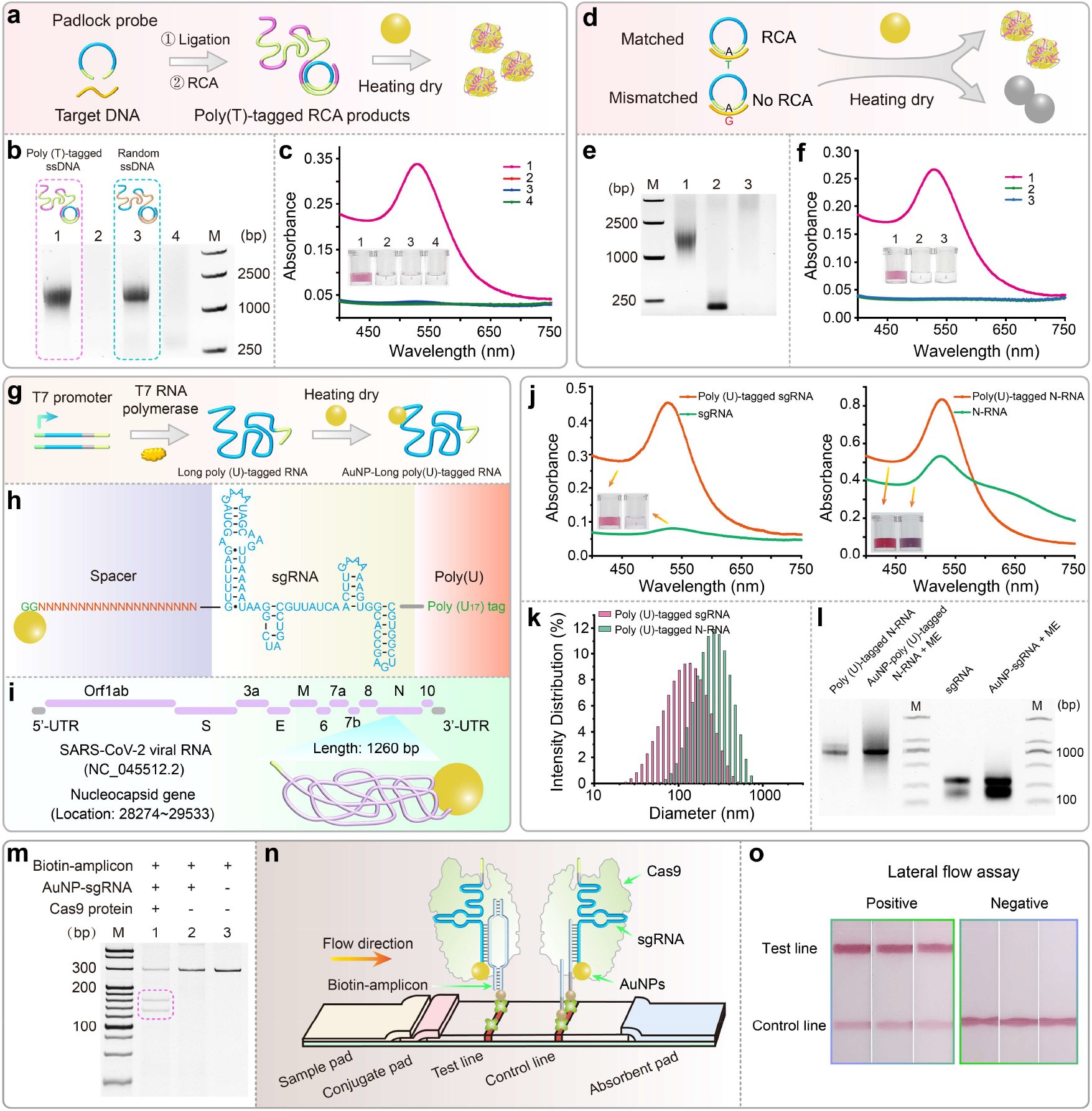
MW-assisted heating-dry method enables labeling of long DNA and long-structured RNA and the application on biosensing. **(a)** MW-assisted heating-dry method for labeling RCA products containing interspaced poly (T) tag. **(b)** Gel electrophoresis analysis of RCA products. Lanes 1 and 3 represent the poly (T)-contained ssDNA and random ssDNA from RCA reaction. Lanes 2 and 4 are the control samples without target DNA template. **(c)** Photographs and the corresponding absorption spectroscopy data showing the labeling results of RCA products. **(d)** Development of colorimetric single-base mutation diagnosis based on ligation-dependent RCA reaction and MW-assisted heating-dry method. **(e)** Gel electrophoresis analysis of RCA products with perfectly matched or single-base mismatched target DNA. Mismatched target DNA will lead to failed ligation and RCA reaction. Lane 1 represents the RCA products with matched DNA, lane 2 represents the RCA reaction products with single mismatched DNA, lane 3 represents the control sample without target DNA. **(f)** Photographs and the corresponding absorption spectroscopy data showing the detection results. **(g)** Labeling of long-structured RNA with poly (U)-tagged sgRNA (135 bp) and long N-RNA (1279 bp) was obtained using *in vitro* T7 transcription. The sequence, structure, and labeling details of sgRNA and N-RNA are shown in **(h)** and **(i)**, respectively. **(j)** Characteristic of labeling results of sgRNA and N-RNA using absorption spectroscopy. **(k)** Dynamic light scattering measurement of the AuNP-sgRNA and AuNP-N-RNA conjugates. **(I)** Parallel gel analysis of the in vitro transcribed sgRNA and N-RNA, and sgRNA and N-RNA that were displaced from the AuNPs surface by ME. **(m)** Electrophoresis analysis of the cleavage ability of the assembled Cas9/sgRNA-AuNP complex. **(n)** Development of CRISPR/Cas9-mediated test strip assay. Schematic illustration of the test strip based on the Cas9/sgRNA-AuNP probes. Cas9/sgRNA-AuNPs probes recognize the biotinylated DNA products and flow through the test strip. The resulting complexes of Cas9/sgRNA-AuNP probe and biotinylated DNA are retained by streptavidin on the test line. Excess Cas9/sgRNA-AuNP probes flow through the control line and are captured by the precoated ssDNA capture probes. **(o)** Photographs taken from Cas9-sgRNA-AuNP probes-based test strip assay for detection of VP72 gene from ASFV.

### MW-assisted heating-dry method enables construction of long and structured ssRNA-AuNP nanostructure

Conjugation of long and structured RNA on AuNPs is of significance due to a wide range of potential applications in biomedicine and nanobiotechnology^46,47^, which, however, remains a challenge. This is because chemical thiol-modification of long-chain RNA is expensive and extremely difficult^25,26^. Currently, high-fidelity chemical synthesis of RNA with more than 200 bases is almost impossible to achieve. Here, we challenged to conjugate AuNPs with long and structured RNA using the heating-dry method. To demonstrate this possibility, poly (U)-tagged sgRNA (135 bp) and Nucleocapsid gene RNA (N-RNA, 1279 bp) of SARS-CoV-2 were obtained using *in vitro* T7 transcription based on a poly (A)-tagged DNA template (Fig. 6g). The sequence, structure, and labeling details of sgRNA and N-RNA are shown in Fig. 6h and i, respectively. sgRNA is a guide RNA in Cas9 gene editing tool and N gene RNA is an important functional gene of SARS-CoV-2. Conjugation of these RNAs to AuNPs may give birth to new medical applications, such as nucleic acid drugs and vaccines^30,31^, delivery of gene editor tool^32,33^, due to the excellent cell entry capabilities of AuNPs^12,48^.

As shown in Fig. 6j, after mixture of RNA and AuNPs and a few minutes of heating-dry process, both the absorption spectra and photographs indicated that poly (U)-tagged sgRNA and N-RNA can be successfully functionalized on AuNPs, while it fails to efficiently load sgRNA and N-RNA without poly (U) tag. The measured hydrodynamic diameters of sgRNA-AuNP and N-RNA-AuNP conjugates are about 141 and 255 nm (Fig. 6k and Supplementary Table 1), respectively, revealing larger RNA strands attached. Next, RNAs were displaced from the AuNPs surface by ME and were subjected to the gel electrophoresis, showing the same map when compared to the original *in vitro* T7 RNA transcription products (Fig. 6l), indicating that RNA sequence was not damaged during the heating-dry process.

Finally, we demonstrated an application of the resulted RNA-AuNP conjugates by developing a CRISPR/Cas9-based nucleic acid test strip using sgRNA-conjugated AuNPs. Previously, we have developed a test strip strategy by employing Cas9/sgRNA as the recognition element to specifically recognize double-stranded DNA (dsDNA), which is superior to the traditional test strip in terms of simplicity, specificity, and portability^49,50^. This conventional Cas9/sgRNA-based test strip employed a DNA-AuNP probe to hybridize with the loop region of sgRNA for lateral flow detection of virus and pathogenic bacteria. Benefiting from the new developed RNA-AuNP conjugation method, now we can further simplify the previous test strip procedure by direct construction of the Cas9/sgRNA-AuNP probes. We next tested if the resulted Cas9/sgRNA-AuNP probes still possess the recognition and cleavage ability. Results showed that both of Cas9/sgRNA and Cas9/sgRNA-AuNP probes could maintain the DNA recognition and cleavage ability (Fig. 6m, n and Supplementary Fig. 16a). To demonstrate the detection application, we amplified the VP72 gene of African swine fever virus (ASFV) with biotinylated primers and designed the corresponding Cas9/sgRNA-AuNP probe to recognize the VP72 gene amplicons (Supplementary Fig. 16b). Subsequent test strip results showed that the positive sample presents a band with high intensity at test line, while the negative sample does not (Fig. 6o), indicating the successful application of the novel Cas9/sgRNA-AuNPs test strip.

## Conclusions

In summary, we invented an ultrafast, general, and cost-effective method to functionalize AuNPs with DNA and RNA using a MW-assisted heating-dry method. The whole labeling process could be completed in minutes by only employing a domestic microwave oven (Supplementary Video). Compared to these state-of-the-art labeling methods, MW-assisted heating-dry method exhibits maximum labeling efficiency, excellent sequence generality, and can produce the most stable DNA-AuNP conjugates (Supplementary Fig. 17). In addition to its confirmed hybridization performance, AuNP-based bioprobes obtained from MW-assisted heating-dry method also exhibit promising new application potential, such as colorimetric DNA single-base mutation analysis and virus DNA detection. Moreover, the heating-dry method has successfully demonstrated its ability for attaching the long-chain and structured nucleic acids, which is difficult to achieve by routine methods. It may stimulate academic interest in branching out into some relatively unexplored areas based on the applications of long-chain DNA/RNA-AuNP conjugates, such as the delivery of gene editing tools, RNA vaccine, and other functional DNA/RNAs.

## Acknowledgments

This work was supported by the National Natural Science Foundation of China (Grants 91959128; 21874049; 21904042; 81772246), Special Project of Science and Technology Development of Guangdong Province (2017B020207011), Guangdong Basic and Applied Basic Research Foundation (2021A1515010262; 2020A1515010754), Special Support Program of Guangdong Province (2016TQ03R749), Opening Project of State Key Laboratory of Chemo/Biosensing and Chemometrics of Hunan University (2019004).

## Author contributions

X.M.Z., E.H.X., and M.Q.H. conceived the study. X.M.Z. and E.H.X. designed and supervised the study. M.Q.H. performed the experiments. M.Q.H. and E.H.X. contributed to the analysis and management of the data. M.Q.H., E.H.X., M.L.H., T.T., H.H.Y., D.B.Z., and X.M.Z. were involved in the data interpretation, discussed the results and commented on the study. X.M.Z., E.H.X., and M.Q.H. wrote the manuscript. All authors reviewed and approved the manuscript.

## Competing interests

The authors declare no competing interests.

## Notes

### Competing Interest Statement

The authors have declared no competing interest.

