## Supplementary Information for "Ultrafast, one-step, and microwave heating-based synthesis of DNA/RNA-AuNP conjugates"

**Materials.**

All the DNA sequences synthesized by Sangon Biotech (Shanghai, China) are listed in Supplementary Table 2. Sodium citrate dihydrate, Sodium hydrogen phosphate, sodium dihydrogen phosphate, 30% acrylamide, ammonium persulfate (APS), N,N,N′,N′-Tetramethylethylenediamine (TEMED), agarose, NaCl, Tris-HCl solution (pH 7.5), ascorbic acid, magnesium chloride, 20× SSC buffer, Tween-20, bovine serum albumin (BSA), n-butanol, HEPES buffer, 20× TAE buffer, 5× TBE buffer, Triton X-100, sucrose, streptavidin, DEPC-treated water, NTPs mix and deoxynucleotide (dNTP) mix were purchased from Sangon Biotech (Shanghai, China). Hexadecyl trimethyl ammonium bromide (CTAB) was purchased from Thermo Fisher Thermo Fisher Scientific. β-mercaptoethanol (ME), sodium borohydride ($NaBH_4$), silver nitrate ($AgNO_3$) and $HAuCl_4$ solution were purchased from Sigma-Aldrich (St. Louis, MO, USA). The Bst 2.0 WarmStart DNA polymerase, DNase I, T4 DNA ligase, and T7 RNA polymerase were supplied by New England Biolabs (Beijing, China). *Streptococcus pyogenes* Cas9 protein was purchased from Bio-lifesci., Ltd. (Guangzhou, China). Premix Ex Taq™ II, 20 bp marker, DL2000 marker, 6× loading-buffer and Recombinant RNase inhibitor (RRI) was purchased from Takara Biotechnology Co., Ltd (Dalian, China). SARS-CoV-2 RNA standards were obtained from National Institute of Metrology (Beijing, China). The RNA Clean & Concentrator Kits was obtained from ZYMO RESEARCH (Beijing, China). The nitrocellulose membrane (NC) was purchased from Sartorius (Shanghai, China). The sample pad, bonding pad, bottom plate and absorbent pad were obtained by Shanghai Jieyi Biotechnology Co., Ltd. (Shanghai, China).

**Apparatus.** The absorption and fluorescence spectra were measured by SpectraMax iD5 Multimode Microplate Reader (Molecular Devices, CA, USA). Real-time fluorescence signal was recorded by a Thermal Cycler Dice™ Real Time System III (Takara, Beijing, China). Domestic microwave oven (MZC-2070M1) that used for heating was purchased from Haier company. The hydrodynamic diameters of particles were measured by a Zetasizer Nano-ZS instrument (Malvern Instruments). The particle size was measured by 200 kV Transmission Electron Microscope (JEM-2100HR, JEOL, Japan). Ultrapure water was provided by Millipore system (Milli-Q, Millipore, >18.25 MΩ·cm). The agarose gel electrophoresis and PAGE experiments were performed using equipment from Beijing Liuyi Instruments (Beijing, China). BG-gdsAUTO320 gel imaging analysis system (Beijing, China) was used for imaging analysis. The XYZ three-dimensional film and spray gold instrument used for preparing lateral flow strip was purchased from Shanghai Jinbiao Biotechnology Co., Ltd. (Shanghai, China). The concentration of nucleic acids was determined by Nanodrop 2000 (Thermo Fisher Scientific). Instrument used for PCR experiment and isothermal incubation were purchased from Eppendorf. Centrifugation of AuNPs probes was performed with Eppendorf centrifuge (5418R, Hamburg, Germany). The Raman Spectroscopy was measured using Renishaw inVia micro-spectrometer (Derbyshire, England). The CD spectrum was collected by a JASCOJ-810 spectropolarimeter (Tokyo, Japan).

**Methods**

**Preparation of 13 nm AuNPs.** The 13 nm AuNPs were synthesized using citrate reduction method as previously reported[1,2]. In brief, 100 mL of 1 mM $HAuCl_4$ solution was added into glass flask and heated to boiling. Then quickly stirred $HAuCl_4$ solution and added 10 mL of 38.8 mM sodium citrate solution. Keep the above solution boiling and stirred for 15~20 min, finally cooled to room temperature with stirring. The prepared AuNPs were stored at 4 °C in the dark.

**Preparation of 30 nm and 40 nm AuNPs.** The 30 nm and 40 nm AuNPs were synthesized using citrate reduction method as previously reported[3]. In brief, 100 mL of 1 mM $HAuCl_4$ solution was added into glass flask and heated to boiling. Then quickly stirred $HAuCl_4$ solution and added 6 mL (5 mL) of 38.8 mM sodium citrate solution for 30 nm (40 nm) AuNPs preparation. Keep the above solution boiling and stirred for 15~20 min, finally cooled to room temperature with stirring. The prepared AuNPs were stored at 4 °C in the dark.

**Preparation of gold nanorods (AuNRs).** The AuNRs were synthesized as previously reported[4]. Firstly, 220 μL of 50 mM $HAuCl_4$ solution was added to 10 mL of 0.2 M CTAB solution, followed by a slight vortex mixing. When the floccule completely dissolved, 21 μL of 0.1 M $AgNO_3$ solution and 212 μL of 0.1 M ascorbic acid were added to the mixture, followed by a slight vortex mixing. Subsequently, 2.5 μL of 1 mM $NaBH_4$ was added to above mixture and incubated at room temperature for 12 h. The prepared AuNRs were washed two times with water and centrifugated (6,000 rpm/min) at 15 °C for 10 min. Finally, the pellet was resuspended in water. The prepared AuNRs were stored at 4 °C in the dark.

**Modification of AuNPs with DNA/RNA using MW-assisted heating-dry method.** 400 μL of AuNPs solution was mixed with 20 μL of 100 μM SH-DNA or poly (T)-DNA or poly (U)-RNA probes in 5 mL glass bottle. Then, put the glass bottle into a domestic microwave oven (the microwave input power is 1150 W and the microwave output power is 700 W) and heated at middle-high model for 3 min, followed by resuspension with $ddH_2O$ or 0.01 M phosphate buffer (0.1 M NaCl, pH 7.4). After that, the particles were washed with 0.01 M phosphate buffer (0.3 M NaCl, pH 7.4) and centrifugated (12,000 rpm/min) at 15 °C for 20 min. Finally, the pellet was resuspended in buffers with different NaCl concentrations, and the resulted nucleic acid-AuNP conjugates were stored at 4 °C in the dark or used for subsequent experiment.

**Modification of AuNPs using salt-aging method.** For salt-aging method[5], 400 μL of AuNPs solution was mixed with 20 μL of 100 μM poly (A/T/C/G)-tagged DNA probes and incubated for 24 h. Then, 40 μL of 125 mM sodium phosphate buffer (pH 7.4) and 40 μL of 1.25 M NaCl was gradually added into the mixture. After incubating for 40 h, the particles were washed three times with 0.01 M phosphate buffer (0.1 M NaCl, pH 7.4) through centrifugation (12000 rpm/min) at 4 °C for 20 min to remove excess DNA strands. Finally, the resulted conjugates were resuspended in buffers with different concentration of NaCl.

**Modification of AuNPs using low-pH method.** For low-pH method[6], 400 μL of AuNPs solution was mixed with 20 μL of 100 μM poly (A/T/C/G)-tagged DNA probes. Then, 8 μL of 500 mM citrate buffer (pH 3) was added to lower the pH of the solution. After brief vortex mixing, the mixture was incubated at room temperature for 3 min. Next, 12 μL of 1 M HEPES buffer (pH 7.6) was added to the mixture to adjust the pH back to a neutral state. Subsequently, the particles were washed three times with 0.01 M phosphate buffer (0.1 M NaCl, pH 7.4) through centrifugation (12000 rpm/min) at 4 °C for 20 min. Finally, the resulted conjugates were resuspended in buffers with different concentrations of NaCl.

**Modification of AuNPs using freeze-thaw method.** For freeze-thaw method[7,8], 400 μL of AuNPs solution was mixed with 20 μL of 100 μM poly (A/T/C/G)-tagged DNA probes. The above mixture was frozen at –20 °C for 1 h and then thawed at room temperature. After that, the particles were washed three times with 0.01 M phosphate buffer (0.1 M NaCl, pH 7.4) through centrifugation (12000 rpm/min) at 4 °C for 20 min. Finally, the resulted conjugates were resuspended in buffers with different concentrations of NaCl.

**Modification of AuNPs using butanol extraction method.** For butanol extraction method[9], 100 μL of AuNPs solution was mixed with 5 μL of 100 μM poly (A/T/C/G)-tagged DNA probes. And then, 900 μL of n-butanol was added to the mixture, followed by a quick vortex mixing. Subsequently, 200 μL of 0.5× TBE (Tris, 44.5 mM; EDTA, 1 mM; boric acid, 44.5 mM; pH 8.0) buffer was added to the above solution followed by quick vortex mixing. The mixture was centrifugated at 2000 g for several seconds to facilitate a liquid phase separation. The liquid phase with particles was pipetted and washed three times with 0.01 M phosphate buffer (0.1 M NaCl, pH 7.4) through centrifugation (12000 rpm/min) at 4 °C for 20 min. Finally, the particles were resuspended in buffers with different concentrations of NaCl.

**Fluorescence quantification of the surface density of DNA-AuNP conjugates.** Poly (T)-tagged DNA used in these studies were labeled with FAM at 3′-terminal (Supplementary Table 2). FAM-labeled DNA were first absorbed to AuNPs surface following the protocol as above described. The FAM-DNA-AuNP conjugates were resuspended in 0.01 M phosphate buffer (0.3 M NaCl, pH 7.4) or 10 mM HEPES buffer. The surface density of the DNA-AuNP conjugate was quantitated according to the published protocol[10]. First, β-mercaptoethanol (ME) was added (final concentration is 20 mM) to the FAM-labeled DNA-AuNPs solution and incubated overnight with shaking at room temperature. Then, the mixture was centrifugated at 15 °C at 12,000 rpm/min for 20 min and the supernatant was collected. The released FAM-DNA probes were in the supernatant and the fluorescence was measured by SpectraMax iD5 Multimode Microplate Reader (Molecular Devices, CA, USA). The fluorescence intensity was converted to the molar concentration of DNA probes by interpolation from a standard linear calibration curve that was prepared with known concentrations of same FAM-DNA probes at same buffer condition and ME concentration. AuNPs concentration was determined via the absorption of UV-vis spectra and finally the number of DNA on each AuNP could be calculated.

**Surface-enhanced Raman scattering (SERS) analysis.** The DNA-AuNP conjugates were obtained using MW-based heating-drying method as above-described and resuspended in 0.01 M phosphate buffer (0.3 M NaCl, pH 7.4). 10 μL of 10 μM ROX-labeled DNA probes complementary to DNA-AuNP conjugates were firstly added to 200 μL DNA-AuNP solution. After 2 h incubation at 37 °C for efficient hybridization, the particles were washed three time with 0.01 M phosphate buffer (0.3 M NaCl, pH 7.4) and centrifugated at 15 °C at 12,000 rpm/min for 20 min. Then the supernatant was discarded and the pellet (20 μL) was collected for SERS measurements. For SERS analysis, the 20 μL pellet was dropped on the Si substrate and the SERS measurement was performed using Renishaw inVia micro-spectrometer (Derbyshire, England). A He–Ne laser operating at $\lambda$ = 633 nm was used as the excitation source with a laser power of approximately 10 mW. All the Raman spectra reported in this work were collected for exposure times of 10 s in the range from $1000^{-1}$ to 1800 $cm^{-1}$.

**Circular dichroism (CD) measurements.** The G4-DNA-AuNP conjugates were obtained using MW-based heating-drying method as above-described. The G4-DNA was diluted in 50 mM Tris-HCl (100 mM KCl, pH 8.5). The G4-DNA-AuNP conjugates were washed two times with 50 mM Tris-HCl (100 mM KCl, pH 8.5) and centrifugated at 15 °C at 12,000 rpm/min for 20 min, followed by resuspending in 50 mM Tris-HCl (100 mM KCl, pH 8.5). The CD spectra of G4-DNA and G4-DNA-AuNP conjugates were collected by a JASCOJ-810 spectropolarimeter (Tokyo, Japan), of which the lamp was always kept under dry purified nitrogen condition during measurements. Three times of scanning from 220 to 310 nm with the scan rate of 100 nm/min were performed and averaged. The background signal of the 50 mM Tris-HCl (100 mM KCl, pH 8.5) buffer was subtracted from the CD data.

**Preparation of the test strip device.** The test trip device consists of sample pad, conjugate pad, nitrocellulose (NC) filter membrane, test line, control line, absorbent pad, and adhesive pad. The sample pad was firstly immersed into a 50 mM Tris-HCl buffer (0.15 M NaCl, 0.25% Triton X-100, pH 8.0) and then dried at 37 °C for 2 h. Streptavidin was embedded in the test line and streptavidin-biotinylated DNA probes was embedded in the control line. The intervals between two adjacent lines were 6 mm. Then the NC membrane was dried at 37 °C for 1 h. Subsequently, the sample pad, conjugate pad, and absorbent pad were attached to the adhesive pad with 2 mm overlap. Finally, the prepared test strip was cut with a width of 4 mm and stored at 4 °C.

**Test strip assay to evaluate the hybridization efficiency of DNA-AuNP conjugates.** The DNA-AuNP conjugates were obtained using MW-assisted heating-dry method. The 200 μL DNA-AuNP solution was washed twice with 0.01 M phosphate buffer (0.1 M NaCl, pH 7.4) by centrifugation at 15 °C at 12,000 rpm/min for 20 min. Subsequently, the pellet was resuspended in 200 μL resuspension buffer (20 mM $Na_3PO_4$, 5% BSA, 0.25% Tween-20, and 10% sucrose). 10 μL resuspended DNA-AuNP conjugates were mixed with 10 μL running buffer (4× SSC, 0.05% Tween-20 (*v*/*v*), 1× PBS, and 1% BSA), and then the above mixture was added to the sample pad of the test trip device. Finally, 100 μL running buffer was added on the sample pad to wash the test strip, and the test strip was photographed after 5 min.

**Synthesis of sgRNA.** sgRNA and poly (U)-tagged sgRNA have been transcribed *in vitro*. The DNA template with or without poly (T) sequence was obtained as previous report by a fill-in PCR method[11]. 500 μL transcription reactions were performed with 0.5 mM NTPs, 250 U T7 RNA polymerase, 50 U RRI, and 500 ng DNA template. The mixture was incubated for 12 h at 37 °C. Then, DNase I was added to digest the DNA template and the RNA transcripts were purified by RNA Clean & Concentrator Kits. The concentration of RNA products was determined by Nanodrop 2000.

**Synthesis of long RNA of Nucleocapsid (N) gene of SARS-CoV-2.** Poly (U)-tagged long RNA (1279 bp) of Nucleocapsid (N) gene of SARS-CoV-2 has been transcribed *in vitro*. The DNA template with or without poly (T) sequence was obtained by a RT-PCR of SARS-CoV-2 RNA standards. The primers were designed according to the sequence of Nucleocapsid (N) gene (Supplementary Table 3). The RT-PCR thermal cycling program: reverser transcription at 50 °C for 45 min, pre-denaturation of cDNA at 94 °C for 2 min, followed by 40 cycles of amplification at 94 °C for 30 s, 55 °C for 30 s, 72 °C for 2 min, finally incubated at 72 °C for 10 min. The PCR products containing T7 promoter was verified by gel electrophoresis and the template was collected using gel extraction kit. The purified DNA templates were then used for *in vitro* transcription system as above described. RNA transcripts were purified and concentrated by RNA Clean & Concentrator Kits and the resulted RNA concentration was determined by Nanodrop 2000.

**Padlock-based ligation and RCA reaction.** Ligation of the linear padlock probe was performed in a 10 μL reaction solution containing 10 mM Tris-HCl buffer (50 mM KCl, 1.5 mM $MgCl_2$, pH 8.9), 500 nM padlock probe, 500 nM matched target DNA, 10 U T4 ligase. The above solution was incubated at 37 °C for 30 min. Subsequently, a total of 20 μL RCA reaction system containing 10 mM Tris-HCl buffer (50 mM KCl, 1.5 mM $MgCl_2$, pH 8.9), 1.25 mM dNTP mix, 3.2 U Bst 2.0 WarmStart DNA polymerase, and 10 μL ligation products, was perform at 60 °C for 60 min. The RCA products were analyzed by gel electrophoresis.

**Loading of AuNPs with RCA products using MW-assisted heating-dry method.** 400 μL of AuNPs solution was mixed with 20 μL RCA products in 5 mL glass bottle. Then, put the glass bottle into a microwave oven and heated at middle-high model for 3 min, followed by resuspension with dd$H_2O$. After that, the solution was washed with 2 mM Tris-HCl buffer (10 mM KCl, 0.3

mM $MgCl_2$, pH 8.9) and then washed with water by centrifugation at 15 °C at 12,000 rpm/min for 20 min. Finally, the pellet was resuspended in 1 M NaCl solution.

**Modification of AuNPs with poly (U)-tagged sgRNA and poly (U)-tagged long RNA of Nucleocapsid (N) gene (1260 bp) of SARS-CoV-2.** 400 μL of AuNPs solution was mixed with 10 μL of 50 μM poly (U)-tagged sgRNA or 320 μg poly (U)-tagged long RNA in 5 mL glass bottle. Then, put the glass bottle into microwave oven and heated at middle-high model for 3min, followed by resuspension with $ddH_2O$. After that, the solution was firstly washed with 0.02 M phosphate buffer (0.3 M NaCl, pH 7.4) and then washed with water by centrifugation at 15 °C at 12,000 rpm/min for 20 min. Finally, the pellet was resuspended in 1 M NaCl solution.

**Test strip assay for nucleic acid detection based on Cas9/sgRNA-AuNP conjugates.** 200 μL sgRNA-AuNP conjugates were firstly washed with $ddH_2O$ for 2 times by centrifugation at 15 °C at 12,000 rpm/min for 20 min and the pellet was resuspended in 10 mM Tris-HCl buffer (50 mM KCl, 1.5 mM $MgCl_2$, pH 8.9). The test strip assay was performed in 20 μL reaction solution included 200 nM Cas9 protein, 15 μL sgRNA-AuNP conjugates, 0.5 μL of 100 mM Tris-HCl buffer (500 mM KCl, 15 mM $MgCl_2$, pH 8.9) and 2 μL PCR products. The mixture was incubated at 37 °C for 30 min. Then, the above solution was added to the sample pad of the test strip device. Subsequently, 50 μL running buffer (4× SSC, 0.05% Tween-20 (*v*/*v*), 1× PBS, and 1% BSA) was added on the sample pad. After 2 min, photographed the lateral flow device.

**Amplification of VP72 Gene of ASFV by PCR.** The PCR reaction of the VP72 gene was performed using a Premix Ex Taq™ II. PCR primers were designed according to the VP72 conserved region of African swine fever virus (ASFV) (Supplementary Table 3). A total of 40 μL amplification system contained 500 nM forward and reverse primer, the genome samples and 1× PCR reagent. The PCR thermal cycling program was: 95 °C for 30 s, followed by 40 cycles of amplification at 95 °C for 5 s, 59 °C for 30 s, a final extension at 72 °C for 10 min and a 10 °C hold. The PCR products were analyzed by gel electrophoresis.

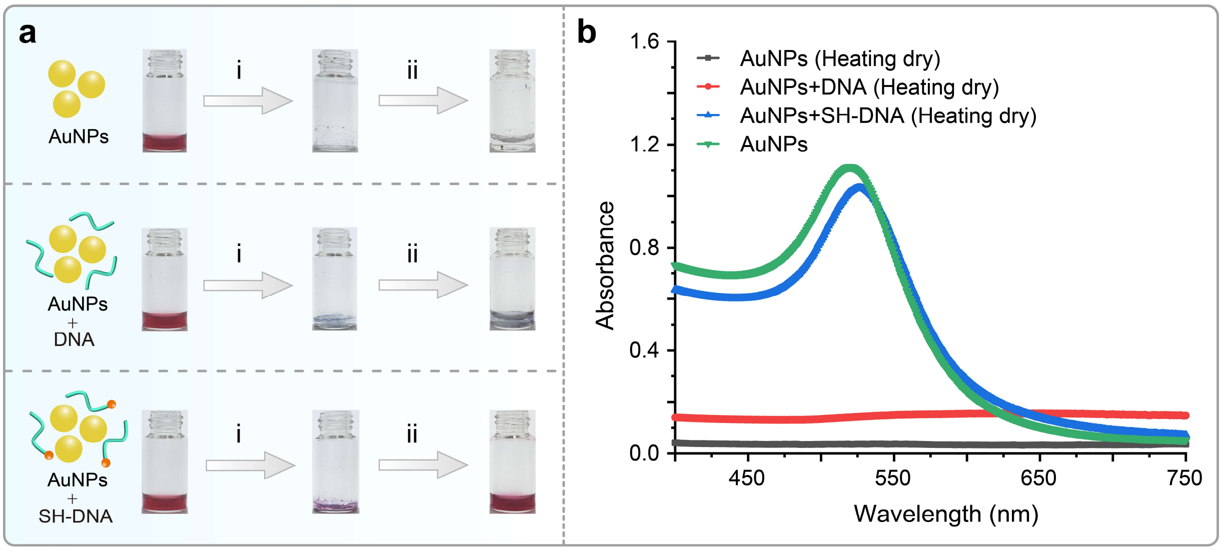


**Supplementary Fig. 1 Ultrafast construction of thiolated DNA/RNA-AuNP conjugates based on MW-assisted heating-dry method. (a)** Photographs of AuNPs before and after heating-dry process. The bare AuNPs and DNA/RNA sequence mixed AuNPs aggregated while SH-DNA/AuNPs retained monodispersed and seems red after heating dry and resuspension. i: MW-assisted heating dry; ii: resuspending with water or buffer. **(b)** Characterization of the SH-DNA-AuNP conjugates based on MW-assisted heating-dry labeling method using UV-vis absorption spectroscopy. The absorption peak of AuNPs with SH-DNA exhibits a 6 nm red-shift, suggesting that there are DNA attached on the AuNPs surface.

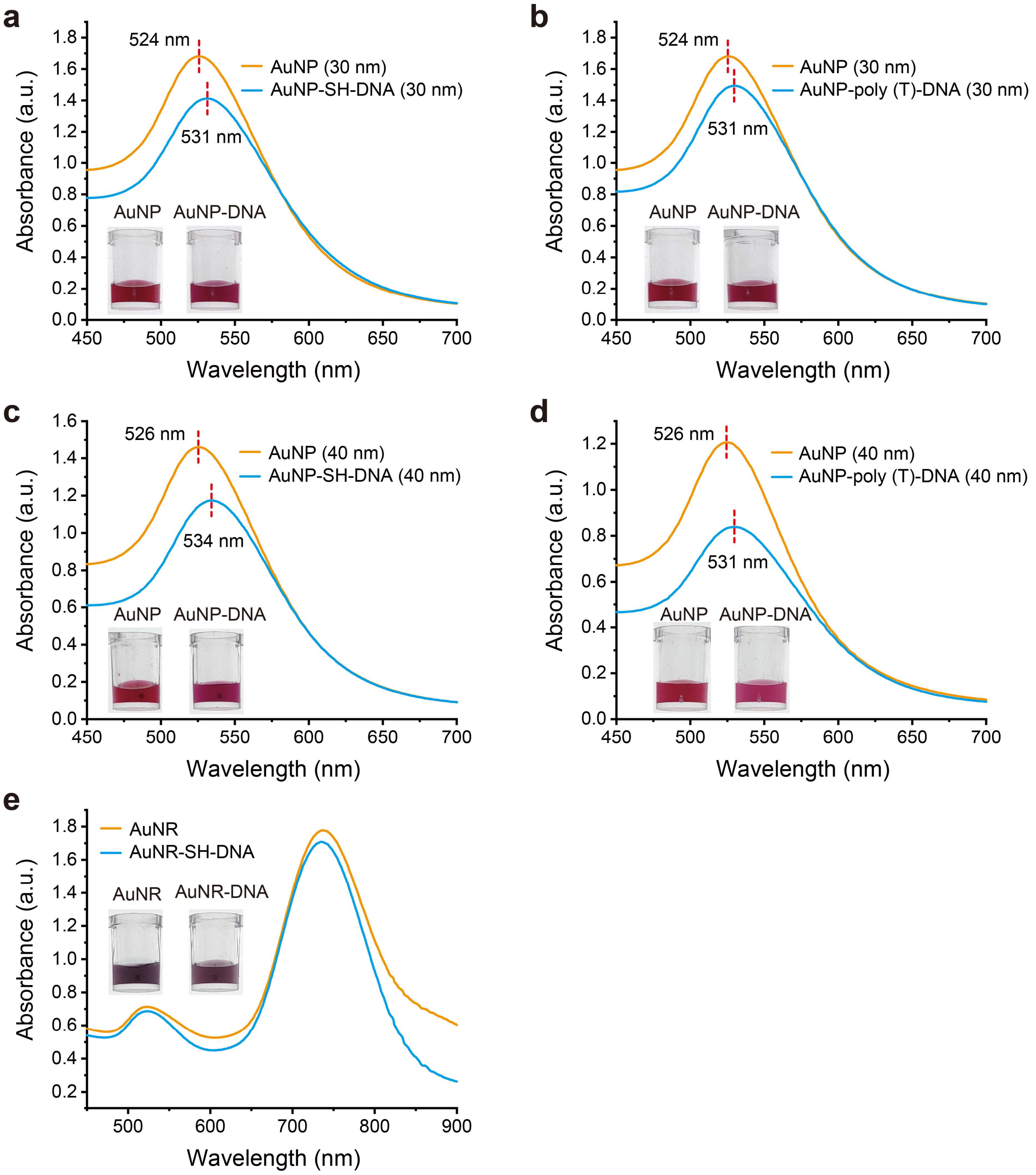


**Supplementary Fig. 2 MW-assisted heating-dry labeling of thiolated and non-thiolated DNA with larger-sized AuNPs and AuNRs. (a)** UV-vis absorption spectroscopy and photographs of 30 nm AuNPs before and after heating-dry labeling of SH-DNA. **(b)** UV-vis absorption spectroscopy and photographs of 30 nm AuNPs before and after heating-dry labeling of poly (T)-tagged DNA. Compared to the bare AuNPs, a 7 nm red-shift is observed, suggesting that SH-DNA and poly (T)-tagged DNA successfully attached on the 30 nm AuNPs surface. **(c)** UV-vis absorption spectroscopy and photographs of 40 nm AuNPs before and after heating-dry labeling of SH-DNA. **(d)** UV-vis absorption spectroscopy and photographs of 40 nm AuNPs before and after heating-dry labeling of poly (T)-tagged DNA. Compared to the bare AuNPs, 8 and 5 nm red-shift are observed, suggesting that SH-DNA and poly (T)-tagged DNA successfully attached on the 40 nm AuNPs surface. **(e)** UV-vis absorption spectroscopy and photographs of AuNRs before and after heating-dry labeling of SH-DNA.

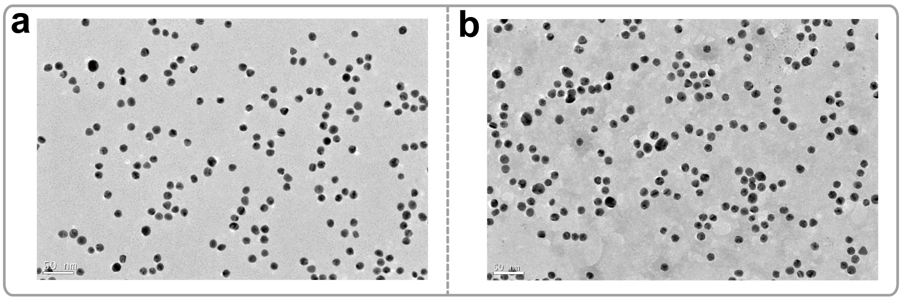


**Supplementary Fig. 3 Transmission electron microscope analysis. (a)** Bare AuNPs, **(b)** Poly (T)-tagged DNA-AuNP conjugates.

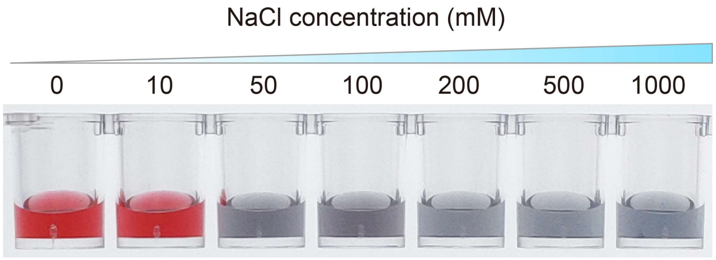


**Supplementary Fig. 4** Salt stability of the bare AuNPs.

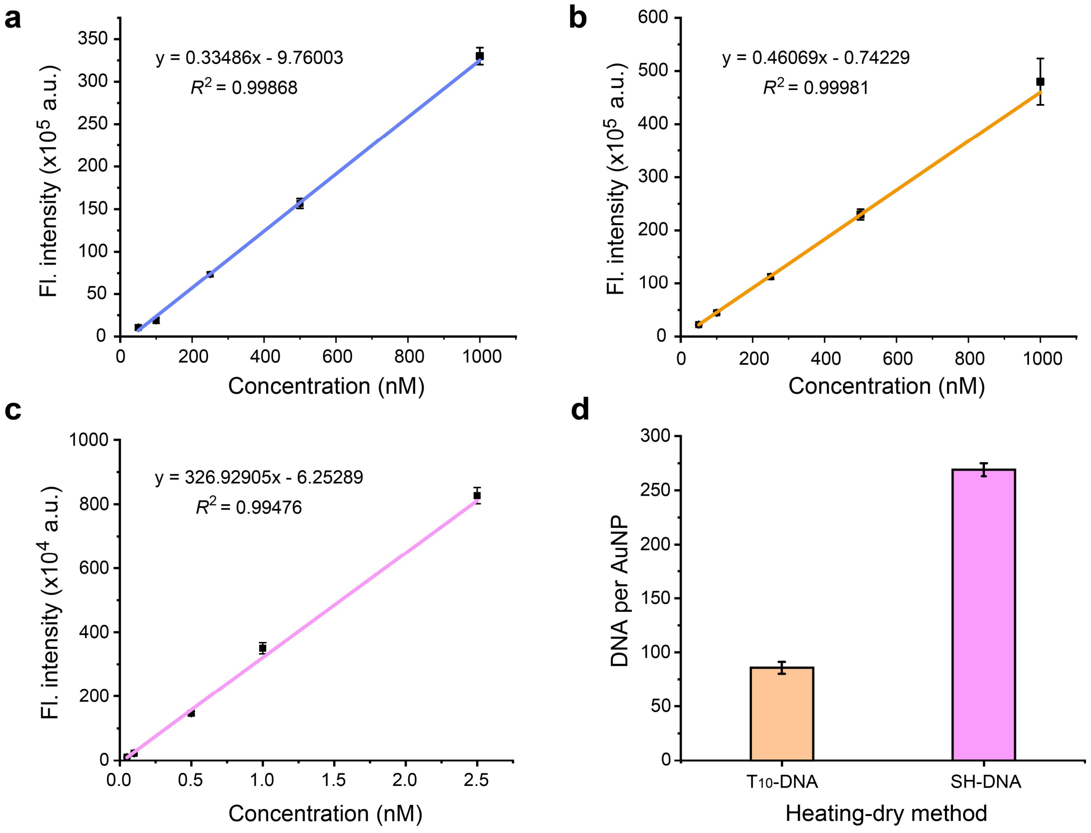


**Supplementary Fig. 5 Analysis the numbers of poly (T)-tagged DNA probes and SH-DNA probes attached to each AuNP. (a)** The fluorescence standard curve of FAM-labelled poly (A)-tagged DNA probe. **(b)** The fluorescence standard curve of FAM-labelled poly (T)-tagged DNA probe. **(c)** The fluorescence standard curve of FAM-labelled SH-DNA probe. **(d)** The measured number of DNA probes attached to each AuNP. The DNA-AuNP conjugates were constructed with poly ($T_{10}$)-tagged DNA and SH-DNA using MW-assisted heating-dry method. Error bars represent three different individual experiments. The number of SH-DNA attached to each AuNP is 269, which is about three times as much as poly (T)-tagged DNA.

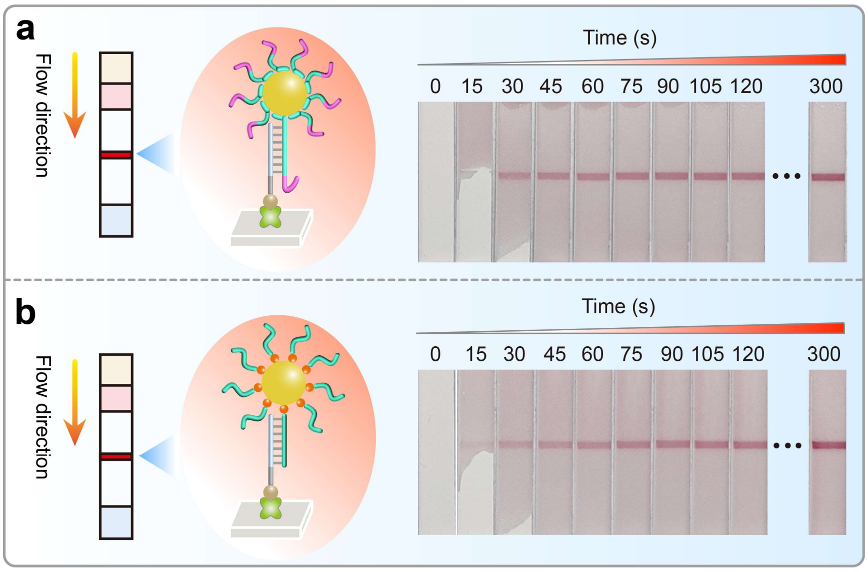


**Supplementary Fig. 6 Evaluation of hybridization ability of non-thiolated and thiolated DNA-AuNP conjugates using test strip assay.** Non-thiolated DNA-AuNPs conjugates were constructed by MW-assisted heating-dry method and SH-DNA-AuNP conjugates were constructed by freeze-thaw method. The streptavidin-biotinylated DNA probes complementary to DNA-AuNP conjugates were pre-embedded in the test strip. For test strip assay, 10 μL resuspended DNA-AuNP conjugates were firstly mixed with 10 μL running buffer and then added to the sample pad of the test strip device. Then, 100 μL running buffer was added on the sample pad to wash the test strip, and the test strip was photographed per 15 s. Photographs showing that the **(a)** non-thiolated as well as **(b)** SH-DNA-AuNP conjugates could be efficiently captured by the biotinylated DNA probes on test strip.

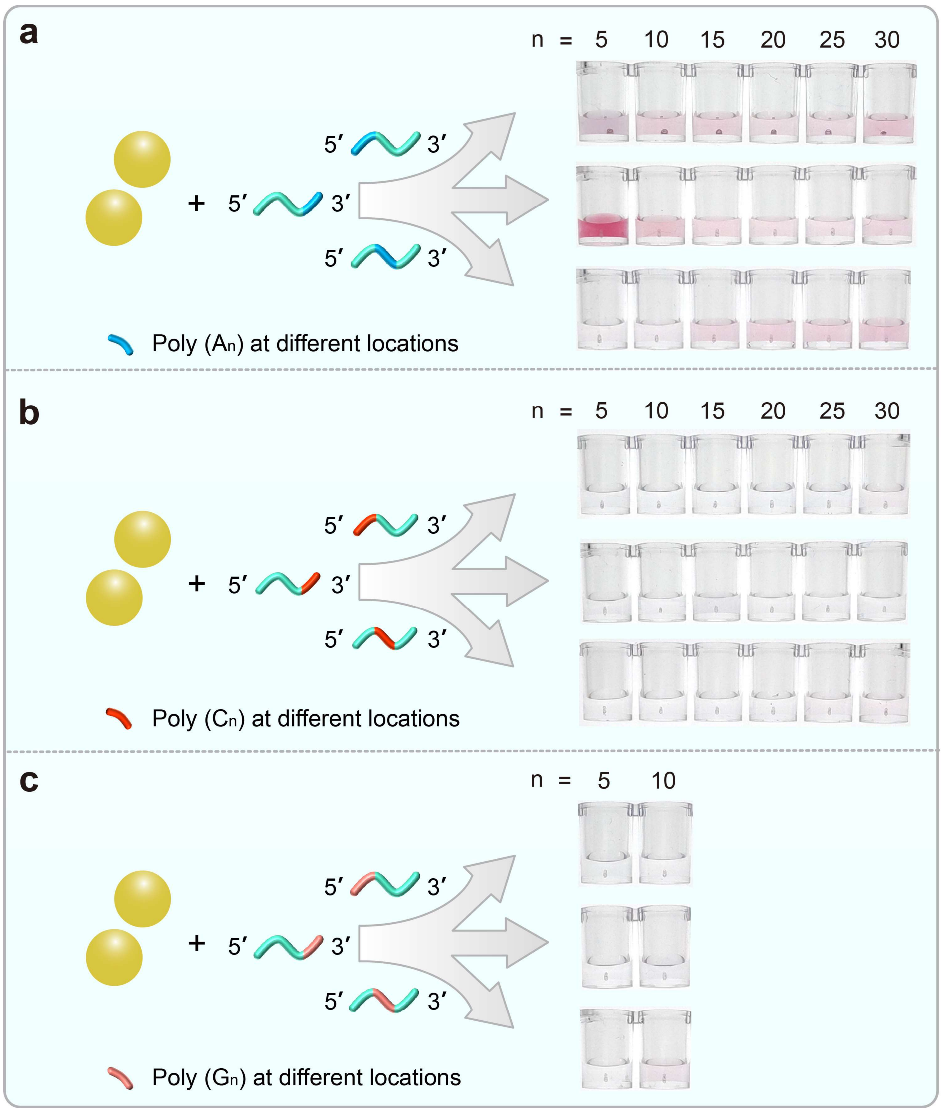


**Supplementary Fig. 7 MW-assisted heating-dry labeling of poly (A/C/G)-tagged DNA. (a)** Photographs showing the labeling results of different lengths of poly (A)-tagged DNA at 5′-terminal, 3′-terminal, and in the middle region. The length of poly (A) tag is 5, 10, 15, 20, 25, and 30, respectively. **(b)** Photographs showing the labeling results of different lengths of poly (C)-tagged DNA at 5′-terminal, 3′-terminal, and in the middle region. The length of poly (A) tag is 5, 10, 15, 20, 25, and 30, respectively. **(c)** Photographs showing the labeling results of different lengths of poly (G)-tagged DNA at 5′-terminal, 3′-terminal, and in the middle region. The length of poly (G) tag is 5 and 10. Longer poly (G) is hard to synthesized. An inefficient and failed labeling will lead to the AuNPs aggregation.

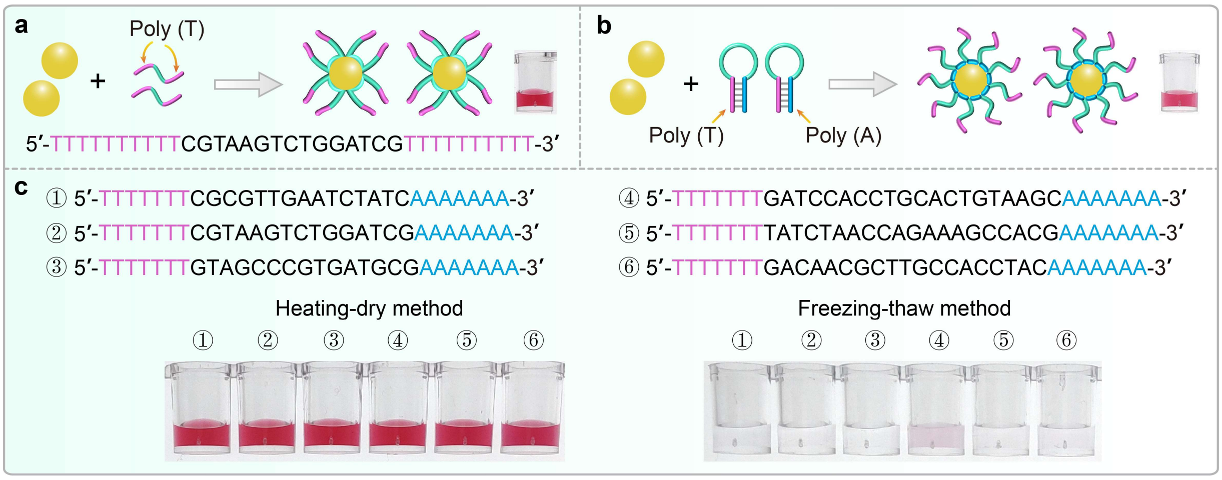


**Supplementary Fig. 8 Labeling of structured DNA. (a)** MW-assisted heating-dry method for labeling of DNA with poly (T) tag located at both 5′- and 3′-terminal. **(b)** MW-assisted heating-dry method for labeling of hairpin DNA sequence with shielded poly (T) tag located in the stem region. **(c)** Photographs showing the labeling results of six hairpin DNA sequences with shielded poly (T) tag located in the stem region using MW-assisted heating-dry method and freeze-thaw method, respectively.

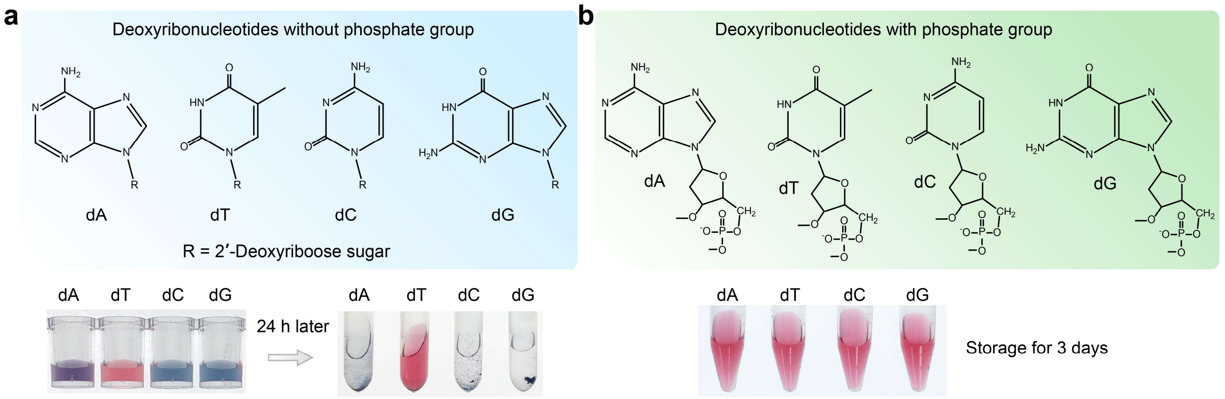


**Supplementary Fig. 9 Colorimetric evaluation of deoxyribonucleotides-Au binding affinity.** **(a)** Photographs showing the binding results of phosphate group-free dA, dT, dC, and dG. 1 μL 5 mM phosphate group-free dA, dT, dC, and dG were mixed with 1 mL 6.4 nM AuNPs solution, respectively. Binding of deoxyribonucleoside to AuNPs resulted in the aggregations of AuNPs due to the replacement of citrate ion, which is a stabilizer of AuNPs. The results showed that phosphate group-free dA, dC, and dG can efficiently bind to AuNP while dT cannot. **(b)** Photographs showing the binding results of phosphate group-contained dA, dT, dC, and dG. 1 μL 5 mM dA, dT, dC, and dG were mixed with 1 mL 6.4 nM AuNPs solution, respectively. Binding of deoxyribonucleotides to AuNPs did not result in the AuNPs aggregations because the surface charge is not replaced.

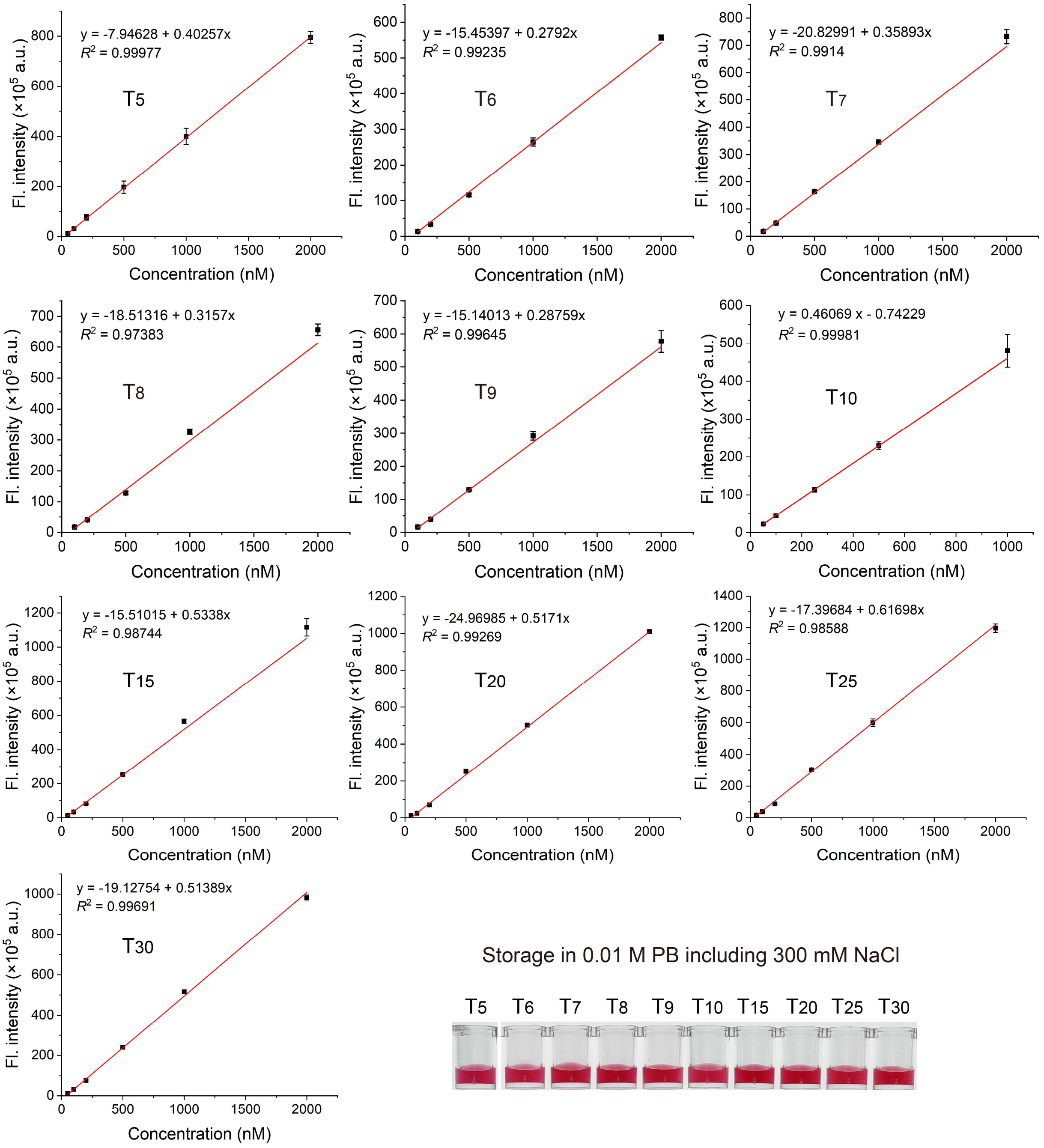


**Supplementary Fig. 10 The fluorescence standard curves of FAM-labelled Poly ($T_n$)-Poly ($C_5$) DNA probes (n = 5, 6, 7, 8, 9, 10, 15, 20, 25, and 30, respectively).** Photographs showing that all Poly ($T_n$)-Poly ($C_5$) DNA probes could be efficiently attached on AuNPs since the AuNPs remains monodisperse and seems red after labeling and resuspending in 0.01 M PB including 300 mM NaCl.

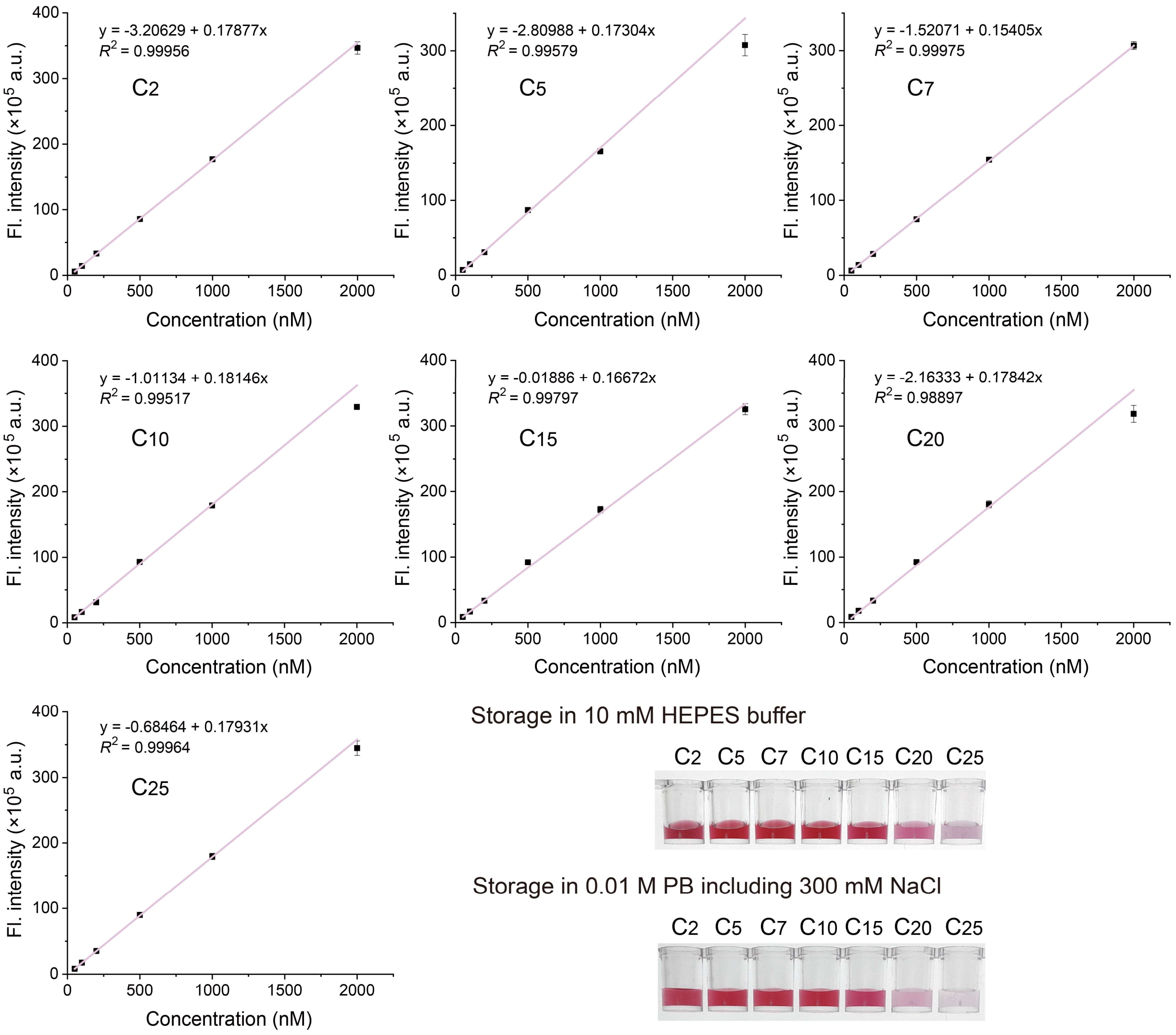


**Supplementary Fig. 11 The fluorescence standard curves of FAM-labelled Poly ($T_{10}$)-Poly ($C_n$) DNA probe (n = 2, 5, 7, 10, 15, 20, and 25, respectively).** Photographs showing the labeling results of these Poly ($T_{10}$)-Poly ($C_n$) DNA. The DNA-AuNP conjugates were resuspended in 10 mM HEPES buffer and 0.01 M PB including 300 mM NaCl. When n is over than 15, the DNA-AuNP aggregated after MW-assisted heating-drying since the number of Poly ($T_{10}$)-Poly ($C_n$) DNA probe attached on each AuNP is not enough to stabilize the AuNPs.

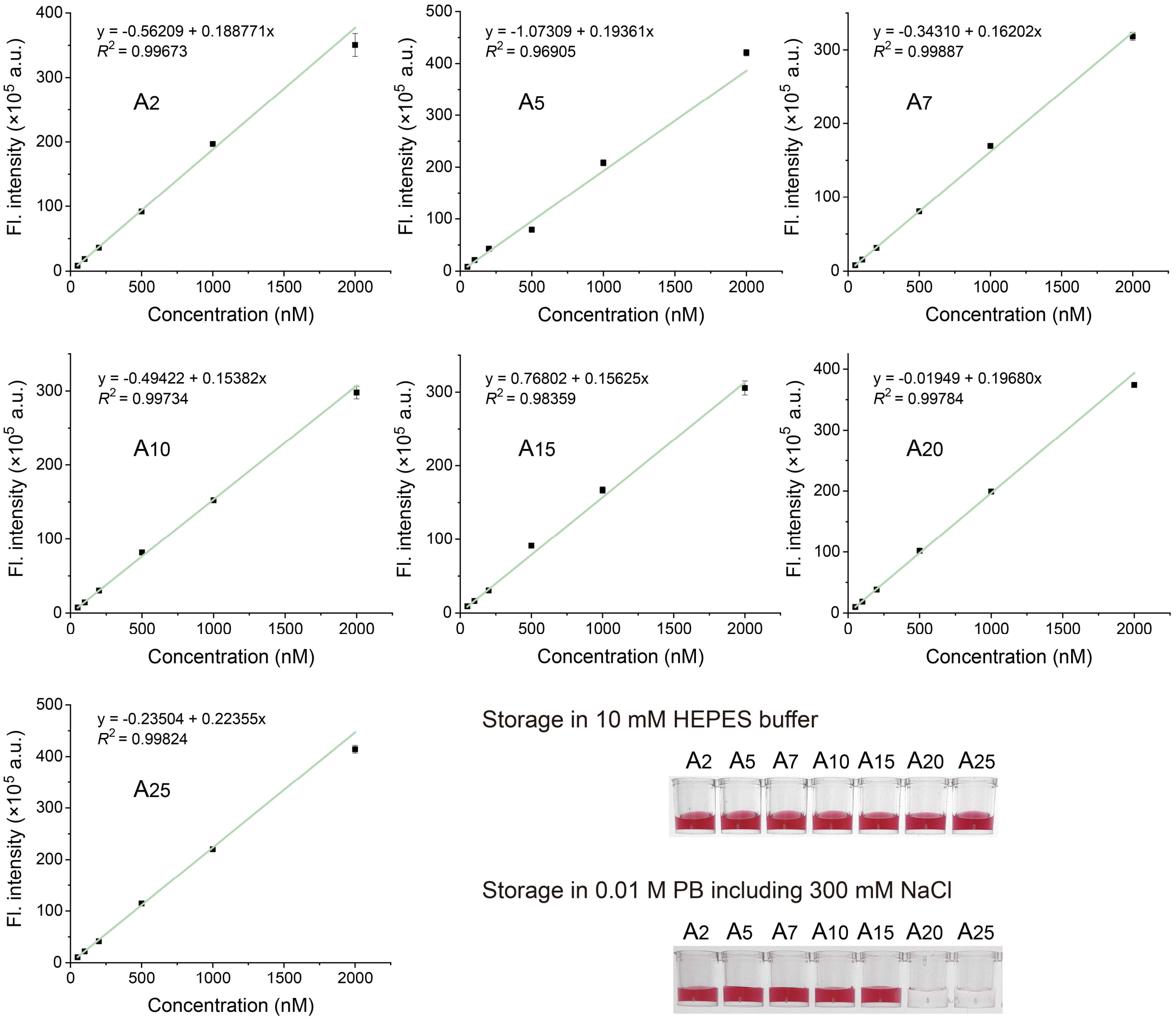


**Supplementary Fig. 12 The fluorescence standard curves of FAM-labelled Poly ($T_{10}$)-Poly (An) DNA probe (n = 2, 5, 7, 10, 15, 20, and 25, respectively).** Photographs showing the labeling results of these Poly ($T_{10}$)-Poly ($A_n$) DNA. The DNA-AuNP conjugates were resuspended in 10 mM HEPES buffer and 0.01 M PB including 300 mM NaCl. When n is over than 15, the DNA-AuNP remains monodisperse and seems red in 10 mM HEPES buffer while aggregated in 0.01 M PB including 300 mM NaCl since the number of Poly ($T_{10}$)-Poly ($A_n$) DNA probe attached on each AuNP is not enough to stabilize the AuNP in 0.01 M PB including 300 mM NaCl.

① 5′-Poly ($A_{10}$/$T_{10}$/$C_{10}$/$G_{10}$)-TATCTAACCAGAAAGCCACG-3′
② 5′-Poly ($A_{10}$/$T_{10}$/$C_{10}$/$G_{10}$)-GACAACGCTTGCCACCTAC-3′
③ 5′-Poly ($A_{10}$/$T_{10}$/$C_{10}$/$G_{10}$)-GATCCACCTGCACTGTAAGC-3′
④ 5′-Poly ($A_{10}$/$T_{10}$/$C_{10}$/$G_{10}$)-CGCGTTGAATCTATC-3′
⑤ 5′-Poly ($A_{10}$/$T_{10}$/$C_{10}$/$G_{10}$)-CGTAAGTCTGGATCG-3′
⑥ 5′-Poly ($A_{10}$/$T_{10}$/$C_{10}$/$G_{10}$)-GTAGCCCGTGATGCG-3′

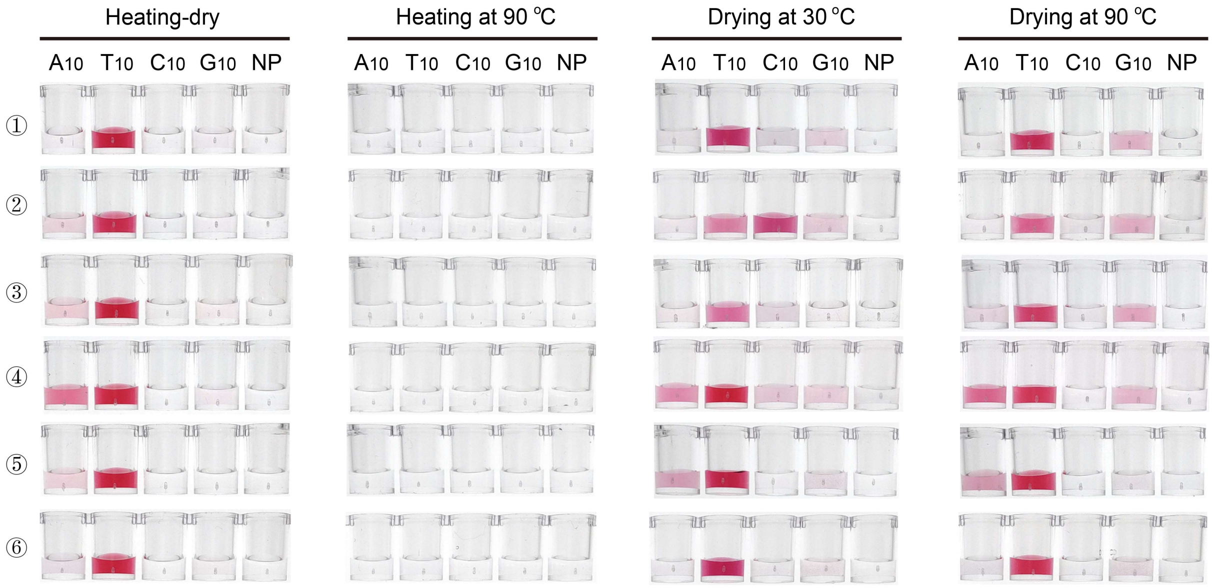


**Supplementary Fig. 13 Photographs showing the labeling results of six poly (A/T/C/G)-tagged DNAs using heating-dry, or heating, or drying method.** The six poly (A/T/C/G)-tagged DNA sequences are shown on the top panel. Heating-drying method was performed as described. For heating method at 90 °C, the same mixture of AuNPs and DNA probes as heating-drying method was heated at 90 °C for 1 h in a sealed tube. For drying method at 30 and 90 °C, the same mixtures of AuNPs and DNA probes as heating-drying method were dried in an air-dry status at 30 °C for 7 h and at 90 °C in an oven-dry status for 1 h, respectively. Photographs showing that all the poly (T)-tagged DNAs exhibited efficient labeling with MW-assisted heating-dry method and failed labeling with heating at 90 °C. For drying method at 30 and 90 °C, the poly (T)-tagged DNA showed decreased and uniform labeling efficiency. While with all the four labeling methods, both the poly (A/C/G)-tagged DNA and non-poly base DNA (NP) exhibited failed labeling.

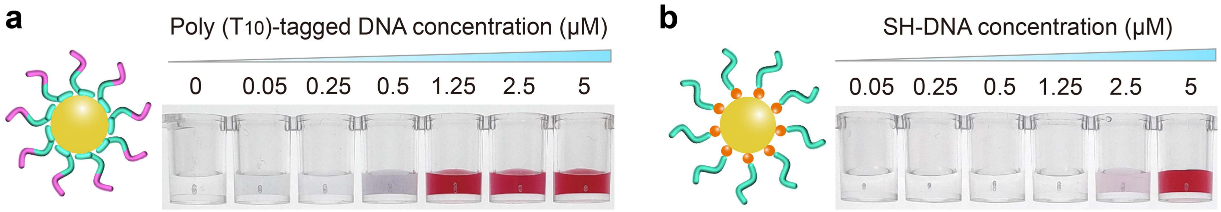


**Supplementary Fig. 14 Test of the critical DNA concentration required for the MW-assisted heating-dry labeling. (a)** Poly ($T_{10}$)-tagged DNA, **(b)** SH-DNA.

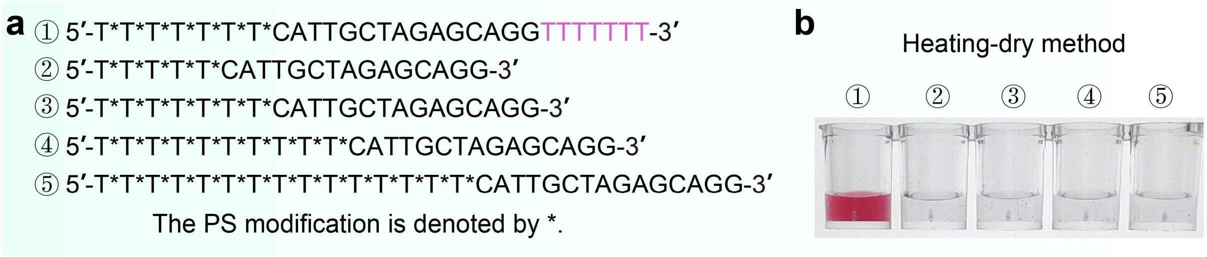


**Supplementary Fig. 15 MW-assisted heating-dry method for labeling of phosphorothioate (PS)-modified DNA sequence. (a)** The detail sequence of PS-modified poly (T)-tagged DNA probes. **(b)** The labeling results of these DNA probes using MW-assisted heating-dry method.

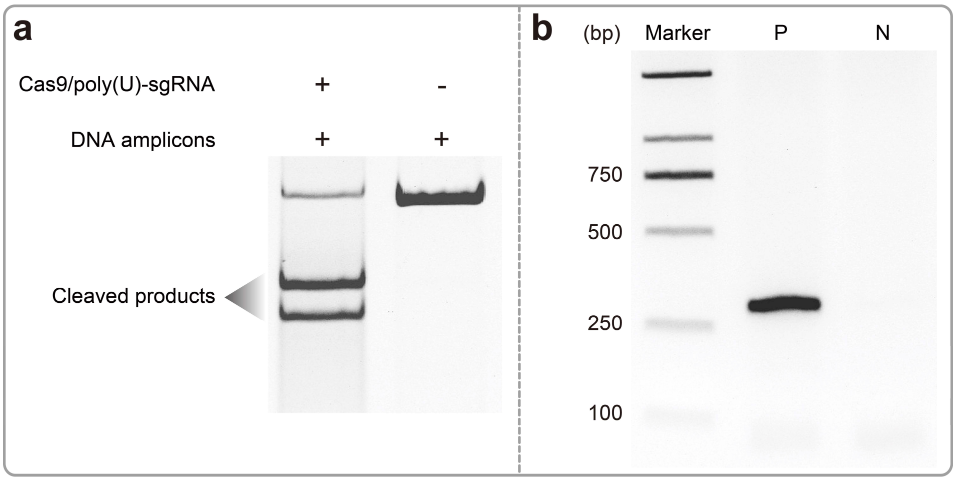


**Supplementary Fig. 16 Gel electrophoretic analysis. (a)** Cleavage ability evaluation using designed poly (U)-tagged sgRNA and Cas9 protein. **(b)** Gel electrophoretic analysis of the PCR products of VP72 gene. P and N stand for positive and negative samples, respectively.

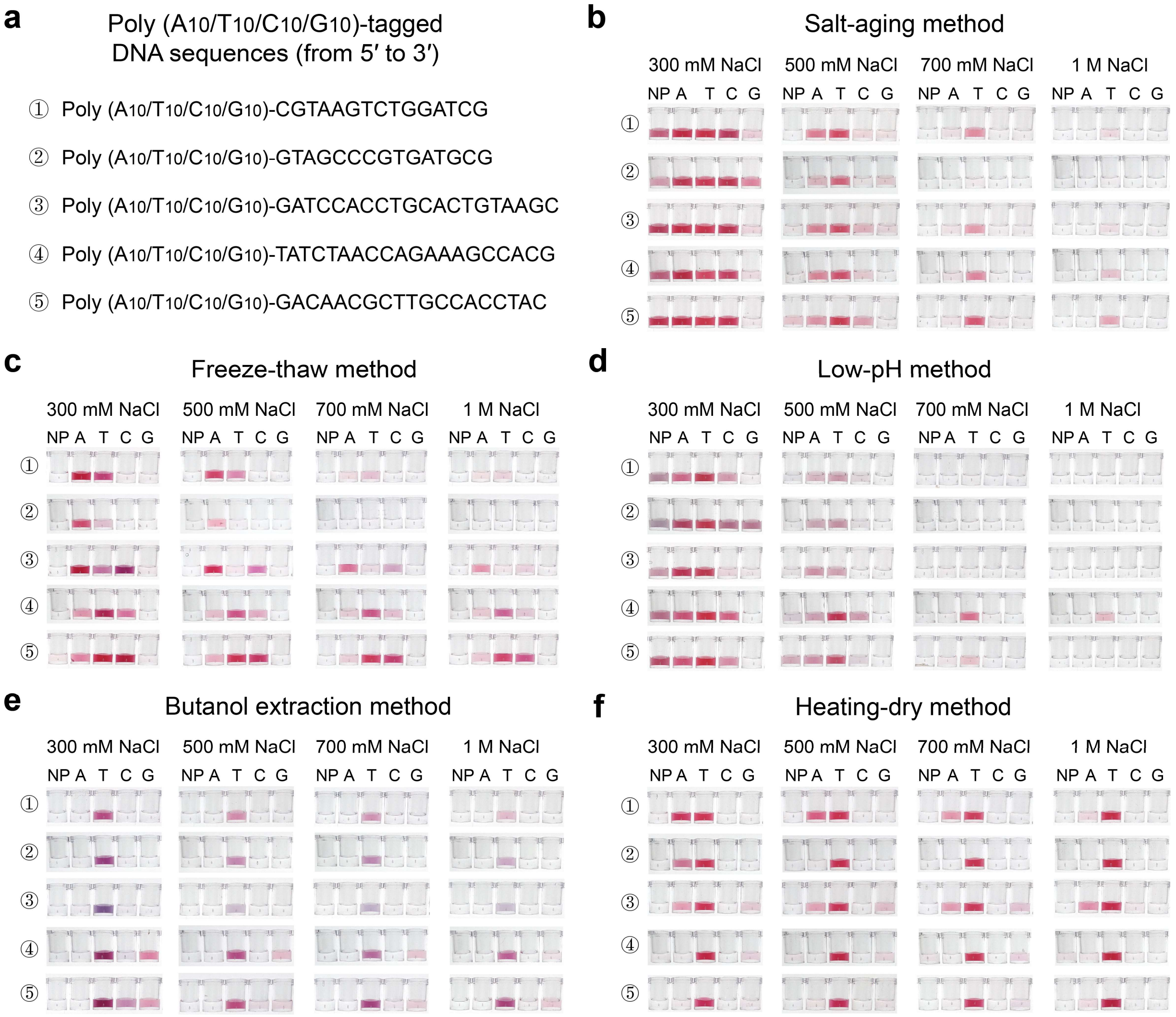


**Supplementary Fig. 17 Non-thiolated DNA-AuNP conjugation with various labeling methods.** **(a)** Detailed sequences of six poly ($A_{10}/T_{10}/C_{10}/G_{10}$)-tagged DNA probes. Photographs showing the labeling results of the six poly ($A_{10}/T_{10}/C_{10}/G_{10}$)-tagged DNA probes using **(b)** salt-aging method. **(c)** freeze-thaw method, **(d)** low-pH method, **(e)** butanol extraction method, and **(f)** MW-assisted heating-dry method. The DNA-AuNP conjugates were resuspended in 300 mM, 500 mM, 700 mM and 1 M NaCl solution, respectively. MW-assisted heating-dry method exhibits maximum labeling efficiency and excellent sequence universality compared with other methods.

**Supplementary Table 1** The mean hydrodynamic diameter and polydispersity index (PDI).

| Names | Average hydrodynamic diameter (nm) | Polydispersity index (PDI) |
|---|---|---|
| **AuNPs** | 22.7 | 0.18 |
| **Poly (T)-tagged DNA-AuNP** | 36.4 | 0.19 |
| **Poly (U)-tagged sgRNA-AuNP** | 161.7 | 0.32 |
| **Poly (U)-tagged N gene-RNA-AuNP** | 285.4 | 0.22 |

**Supplementary Table 2** DNA and RNA sequences used in this work.[a]

| Name | Sequences (from 5′ to 3′) | Locations |
|---|---|---|
| DNA | CGTAAGTCTGGATCG | Fig. 1b, Supplementary Fig. 1a |
| Poly ($T_{10}$)-DNA (5′) | TTTTTTTTTTCGTAAGTCTGGATCG | Fig. 1b-e, j, Fig. 2b, Fig. 5a, Supplementary Figs. 13, 14a |
| FAM-poly ($T_{10}$)-DNA | TTTTTTTTTTCCCCC-FAM | Fig. 1f-i, Supplementary Figs. 2a-e, 5a, 10 |
| FAM-poly ($A_{10}$)-DNA | AAAAAAAAAATTTTT-FAM | Fig. 1f, Supplementary Fig. 5b |
| Poly ($rU_{10}$)-RNA | UUUUUUUUUUCGUAAGUCUGGAUCG | Fig. 1j, Fig. 2a |
| Poly($A_{10}$)-DNA (5′) | AAAAAAAAAACGTAAGTCTGGATCG | Fig. 2a, Fig. 5a, Supplementary Figs. 13, 17 |
| Poly ($C_{10}$)-DNA (5′) | CCCCCCCCCCCGTAAGTCTGGATCG | Fig. 2a, Fig. 5a, Supplementary Fig. 13 |
| Poly ($G_{10}$)-DNA (5′) | GGGGGGGGGGCGTAAGTCTGGATCG | Fig. 2a, Fig. 5a, Supplementary Fig. 13 |
| Poly ($A_{10}$)-DNA (3′) | GCTAGGTCTGAATGCAAAAAAAAAA | Fig. 2a |
| Poly ($T_{10}$)-DNA (3′) | GCTAGGTCTGAATGCTTTTTTTTTT | Fig. 2a, 2b |
| Poly ($C_{10}$)-DNA (3′) | GCTAGGTCTGAATGCCCCCCCCCCC | Fig. 2a |
| Poly ($G_{10}$)-DNA (3′) | GCTAGGTCTGAATGCGGGGGGGGGG | Fig. 2a |
| Poly ($T_{10}$)-DNA-M | CGCGTTGAATCTATCTTTTTTTTTTCGTAAGTCTGGATCG | Fig. 2a, b |
| Poly ($A_{10}$)-DNA-M | CGCGTTGAATCTATCAAAAAAAAAACGTAAGTCTGGATCG | Fig. 2a |
| Poly ($C_{10}$)-DNA-M | CGCGTTGAATCTATCCCCCCCCCCCCGTAAGTCTGGATCG | Fig. 2a |
| Poly ($G_{10}$)-DNA-M | CGCGTTGAATCTATCGGGGGGGGGGCGTAAGTCTGGATCG | Fig. 2a |
| RNA | CGUAAGUCUGGAUCG | Fig. 2a |
| Poly ($rA_{10}$)-RNA | AAAAAAAAAACGUAAGUCUGGAUCG | Fig. 2a |
| Poly ($rC_{10}$)-RNA | CCCCCCCCCCCGUAAGUCUGGAUCG | Fig. 2a |
| Poly ($rG_{10}$)-RNA | GGGGGGGGGGCGUAAGUCUGGAUCG | Fig. 2a |
| Poly ($T_5$)-DNA (5′) | TTTTTCGTAAGTCTGGATCG | Fig. 2b, c |
| Poly ($T_{15}$)-DNA (5′) | TTTTTTTTTTTTTTTCGTAAGTCTGGATCG | Fig. 2b |
| Poly ($T_{20}$)-DNA (5′) | TTTTTTTTTTTTTTTTTTTTCGTAAGTCTGGATCG | Fig. 2b |
| Poly ($T_{25}$)-DNA (5′) | TTTTTTTTTTTTTTTTTTTTTTTTTCGTAAGTCTGGATCG | Fig. 2b |
| Poly ($T_{30}$)-DNA (5′) | TTTTTTTTTTTTTTTTTTTTTTTTTTTTTTCGTAAGTCTGGATCG | Fig. 2b |
| Poly ($T_5$)-DNA (3′) | GCTAGGTCTGAATGCTTTTT | Fig. 2b, c |
| Poly ($T_{15}$)-DNA (3′) | GCTAGGTCTGAATGCTTTTTTTTTTTTTTT | Fig. 2b |
| Poly ($T_{20}$)-DNA (3′) | GCTAGGTCTGAATGCTTTTTTTTTTTTTTTTTTTT | Fig. 2b |

| | | |
|---|---|---|
| Poly ($T_{25}$)-DNA (3′) | GCTAGGTCTGAATGCTTTTTTTTTTTTTTTTT<br>TTTTTTTT | Fig. 2b |
| Poly ($T_{30}$)-DNA (3′) | GCTAGGTCTGAATGCTTTTTTTTTTTTTTTTT<br>TTTTTTTTTTTTT | Fig. 2b |
| Poly ($T_5$)-DNA-M | CGCGTTGAATCTATCTTTTTCGTAAGTCTGG<br>ATCG | Fig. 2b |
| Poly ($T_{15}$)-DNA-M | CGCGTTGAATCTATCTTTTTTTTTTTTTTTCG<br>TAAGTCTGGATCG | Fig. 2b |
| Poly ($T_{20}$)-DNA-M | CGCGTTGAATCTATCTTTTTTTTTTTTTTTTT<br>TTTCGTAAGTCTGGATCG | Fig. 2b |
| Poly ($T_{25}$)-DNA-M | CGCGTTGAATCTATCTTTTTTTTTTTTTTTTT<br>TTTTTTTTCGTAAGTCTGGATCG | Fig. 2b |
| Poly ($T_{30}$)-DNA-M | CGCGTTGAATCTATCTTTTTTTTTTTTTTTTT<br>TTTTTTTTTTTTTCGTAAGTCTGGATCG | Fig. 2b |
| $T_3C_2$-1-DNA (5′) | CCTTTCGTAAGTCTGGATCG | Fig. 2c |
| $T_3C_2$-2-DNA (5′) | TCCTTCGTAAGTCTGGATCG | Fig. 2c |
| $T_3C_2$-3-DNA (5′) | TTTCCCGTAAGTCTGGATCG | Fig. 2c |
| $T_4C_1$-1-DNA (5′) | CTTTTCGTAAGTCTGGATCG | Fig. 2c |
| $T_4C_1$-2-DNA (5′) | TCTTTCGTAAGTCTGGATCG | Fig. 2c |
| $T_4C_1$-3-DNA (5′) | TTCTTCGTAAGTCTGGATCG | Fig. 2c |
| $T_4C_1$-4-DNA (5′) | TTTCTCGTAAGTCTGGATCG | Fig. 2c |
| $T_4C_1$-5-DNA (5′) | TTTTCCGTAAGTCTGGATCG | Fig. 2c |
| $T_3C_2$-1-DNA (3′) | GCTAGGTCTGAATGCTTTCC | Fig. 2c |
| $T_3C_2$-2-DNA (3′) | GCTAGGTCTGAATGCTTCCT | Fig. 2c |
| $T_3C_2$-3-DNA (3′) | GCTAGGTCTGAATGCCCTTT | Fig. 2c |
| $T_4C_1$-1-DNA (3′) | GCTAGGTCTGAATGCTTTTC | Fig. 2c |
| $T_4C_1$-2-DNA (3′) | GCTAGGTCTGAATGCTTTCT | Fig. 2c |
| $T_4C_1$-3-DNA (3′) | GCTAGGTCTGAATGCTTCTT | Fig. 2c |
| $T_4C_1$-4-DNA (3′) | GCTAGGTCTGAATGCTCTTT | Fig. 2c |
| $T_4C_1$-5-DNA (3′) | GCTAGGTCTGAATGCCTTTT | Fig. 2c |
| $T_3A_2$-1-DNA (5′) | AATTTCGTAAGTCTGGATCG | Fig. 2c |
| $T_3A_2$-2-DNA (5′) | TAATTCGTAAGTCTGGATCG | Fig. 2c |
| $T_3A_2$-3-DNA (5′) | TTTAACGTAAGTCTGGATCG | Fig. 2c |
| $T_4A_1$-1-DNA (5′) | ATTTTCGTAAGTCTGGATCG | Fig. 2c |
| $T_4A_1$-2-DNA (5′) | TATTTCGTAAGTCTGGATCG | Fig. 2c |
| $T_4A_1$-3-DNA (5′) | TTATTCGTAAGTCTGGATCG | Fig. 2c |
| $T_4A_1$-4-DNA (5′) | TTTATCGTAAGTCTGGATCG | Fig. 2c |
| $T_4A_1$-5-DNA (5′) | TTTTACGTAAGTCTGGATCG | Fig. 2c |
| $T_3A_2$-1-DNA (3′) | GCTAGGTCTGAATGCTTTAA | Fig. 2c |
| $T_3A_2$-2-DNA (3′) | GCTAGGTCTGAATGCTTAAT | Fig. 2c |
| $T_3A_2$-3-DNA (3′) | GCTAGGTCTGAATGCAATTT | Fig. 2c |
| $T_4A_1$-1-DNA (3′) | GCTAGGTCTGAATGCTTTTA | Fig. 2c |
| $T_4A_1$-2-DNA (3′) | GCTAGGTCTGAATGCTTTAT | Fig. 2c |
| $T_4A_1$-3-DNA (3′) | GCTAGGTCTGAATGCTTATT | Fig. 2c |
| $T_4A_1$-4-DNA (3′) | GCTAGGTCTGAATGCTATTT | Fig. 2c |
| $T_4A_1$-5-DNA (3′) | GCTAGGTCTGAATGCATTTT | Fig. 2c |
| $T_3G_2$-1-DNA (5′) | GGTTTCGTAAGTCTGGATCG | Fig. 2c |
| $T_3G_2$-2-DNA (5′) | TGGTTCGTAAGTCTGGATCG | Fig. 2c |
| $T_3G_2$-3-DNA (5′) | TTTGGCGTAAGTCTGGATCG | Fig. 2c |
| $T_4G_1$-1-DNA (5′) | GTTTTCGTAAGTCTGGATCG | Fig. 2c |

| | | |
|---|---|---|
| $T_4G_1$-2-DNA (5′) | TGTTTCGTAAGTCTGGATCG | Fig. 2c |
| $T_4G_1$-3-DNA (5′) | TTGTTCGTAAGTCTGGATCG | Fig. 2c |
| $T_4G_1$-4-DNA (5′) | TTTGTCGTAAGTCTGGATCG | Fig. 2c |
| $T_4G_1$-5-DNA (5′) | TTTTGCGTAAGTCTGGATCG | Fig. 2c |
| $T_3G_2$-1-DNA (3′) | GCTAGGTCTGAATGCTTTGG | Fig. 2c |
| $T_3G_2$-2-DNA (3′) | GCTAGGTCTGAATGCTTGGT | Fig. 2c |
| $T_3G_2$-3-DNA (3′) | GCTAGGTCTGAATGCGGTTT | Fig. 2c |
| $T_4G_1$-1-DNA (3′) | GCTAGGTCTGAATGCTTTTG | Fig. 2c |
| $T_4G_1$-2-DNA (3′) | GCTAGGTCTGAATGCTTTGT | Fig. 2c |
| $T_4G_1$-3-DNA (3′) | GCTAGGTCTGAATGCTTGTT | Fig. 2c |
| $T_4G_1$-4-DNA (3′) | GCTAGGTCTGAATGCTGTTT | Fig. 2c |
| $T_4C_1$-5-DNA (3′) | GCTAGGTCTGAATGCGTTTT | Fig. 2c |
| $T_{10}A_1$ | TTTTTTTTTTA | Fig. 2d |
| $T_{10}T_1$ | TTTTTTTTTTT | Fig. 2d |
| $T_{10}C_1$ | TTTTTTTTTTC | Fig. 2d |
| $T_{10}G_1$ | TTTTTTTTTTG | Fig. 2d |
| $T_{10}A_2$ | TTTTTTTTTTAA | Fig. 2d |
| $T_{10}T_2$ | TTTTTTTTTTTT | Fig. 2d |
| $T_{10}C_2$ | TTTTTTTTTTCC | Fig. 2d |
| $T_{10}G_2$ | TTTTTTTTTTGG | Fig. 2d |
| $A_{10}A_2$ | AAAAAAAAAAAA | Fig. 2d |
| $A_{10}T_2$ | AAAAAAAAAATT | Fig. 2d |
| $A_{10}C_2$ | AAAAAAAAAACC | Fig. 2d |
| $A_{10}G_2$ | AAAAAAAAAAGG | Fig. 2d |
| $C_{10}A_2$ | CCCCCCCCCCAA | Fig. 2d |
| $C_{10}T_2$ | CCCCCCCCCCTT | Fig. 2d |
| $C_{10}C_2$ | CCCCCCCCCCCC | Fig. 2d |
| $C_{10}G_2$ | CCCCCCCCCCGG | Fig. 2d |
| $G_{10}A_2$ | GGGGGGGGGGAA | Fig. 2d |
| $G_{10}T_2$ | GGGGGGGGGGTT | Fig. 2d |
| $G_{10}C_2$ | GGGGGGGGGGCC | Fig. 2d |
| $G_{10}G_2$ | GGGGGGGGGGGG | Fig. 2d |
| SH-DNA | SH-GCTTACAGTGCAGGTGGATCCGATAC AGACTTACG | Fig. 3a |
| $A_5$-DNA-$T_5$ | AAAAAGCTTACAGTGCAGGTGGATCCGATA CAGACTTACGTTTTT | Fig. 3a |
| $A_5$-DNA-$T_{10}$ | AAAAAGCTTACAGTGCAGGTGGATCCGATA CAGACTTACGTTTTTTTTTT | Fig. 3a |
| ROX-cDNA | ROX-CGTAAGTCTGTATCGGATCCACCTG CACTGTAAGC | Fig. 3a |
| $T_{10}$-G4-DNA1 | TTTTTTTTTTGGTGGTGGTGGTGGTGGTGGT | Fig. 3b |
| $T_{10}$-G4-DNA1-$T_{10}$ | TTTTTTTTTTGGTGGTGGTGGTGGTGGTGGT TTTTTTTTT | Fig. 3b |

| | | |
|---|---|---|
| $T_{10}$-G4-DNA1-$A_5$ | TTTTTTTTTTGGTGGTGGTGGTGGTGGTGGTTTAAAAA | Fig. 3b |
| $T_{10}$-G4-DNA1-$A_{10}$ | TTTTTTTTTTGGTGGTGGTGGTGGTGGTGGTTTAAAAAAAAAA | Fig. 3b |
| $T_{10}$-G4-DNA1-$A_{15}$ | TTTTTTTTTTGGTGGTGGTGGTGGTGGTGGTTTAAAAAAAAAAAAAAA | Fig. 3b |
| $T_{10}$-G4-DNA2 | TTTTTTTTTTGGGTAGGGCGGGTTGGG | Fig. 3c |
| $T_{10}$-G4-DNA2-$T_{10}$ | TTTTTTTTTTGGGTAGGGCGGGTTGGGTTTTTTTTTT | Fig. 3c |
| $T_{10}$-G4-DNA2-$A_5$ | TTTTTTTTTTGGGTAGGGCGGGTTGGGTTTAAAAA | Fig. 3c |
| $T_{10}$-G4-DNA2-$A_{10}$ | TTTTTTTTTTGGGTAGGGCGGGTTGGGTTTAAAAAAAAAA | Fig. 3c |
| $T_{10}$-G4-DNA2-$A_{15}$ | TTTTTTTTTTGGGTAGGGCGGGTTGGGTTTAAAAAAAAAAAAAAA | Fig. 3c |
| Poly ($T_5$)-DNA (hybrid) | TTTTTCATTGCTAGAGCAGG | Fig. 4a |
| Poly ($T_{10}$)-DNA (hybrid) | TTTTTTTTTTCATTGCTAGAGCAGG | Fig. 4a |
| Poly ($T_{15}$)-DNA (hybrid) | TTTTTTTTTTTTTTTCATTGCTAGAGCAGG | Fig. 4a |
| Poly ($T_7$)-DNA-poly($A_0$) (hybrid) | TTTTTTTCATTGCTAGAGCAGG | Fig. 4a |
| Poly ($T_7$)-DNA-poly($A_2$) (hybrid) | TTTTTTTCATTGCTAGAGCAGGAA | Fig. 4a |
| Poly ($T_7$)-DNA-poly($A_5$) (hybrid) | TTTTTTTCATTGCTAGAGCAGGAAAAA | Fig. 4a |
| Poly ($T_7$)-DNA-poly($A_{10}$) (hybrid) | TTTTTTTCATTGCTAGAGCAGGAAAAAAAAAA | Fig. 4a |
| Poly ($T_7$)-DNA-poly($A_{15}$) (hybrid) | TTTTTTTCATTGCTAGAGCAGGAAAAAAAAAAAAAAA | Fig. 4a |
| Poly ($T_7$)-DNA-poly($A_{20}$) (hybrid) | TTTTTTTCATTGCTAGAGCAGGAAAAAAAAAAAAAAAAAAAA | Fig. 4a |
| Poly ($T_7$)-DNA-poly($A_{25}$) (hybrid) | TTTTTTTCATTGCTAGAGCAGGAAAAAAAAAAAAAAAAAAAAAAAAA | Fig. 4a |
| Poly ($T_5$)-Poly($C_5$)-FAM | TTTTTCCCCC-FAM | Fig. 4b, Supplementary Fig. 10 |
| Poly ($T_6$)-Poly($C_5$)-FAM | TTTTTTCCCCC-FAM | Fig. 4b, Supplementary Fig. 10 |
| Poly ($T_7$)-Poly($C_5$)-FAM | TTTTTTTCCCCC-FAM | Fig. 4b, Supplementary Fig. 10 |
| Poly ($T_8$)-Poly($C_5$)-FAM | TTTTTTTTCCCCC-FAM | Fig. 4b, Supplementary Fig. 10 |
| Poly ($T_9$)-Poly($C_5$)-FAM | TTTTTTTTTCCCCC-FAM | Fig. 4b, Supplementary Fig. 10 |
| Poly ($T_{15}$)-Poly ($C_5$)-FAM | TTTTTTTTTTTTTTTCCCCC-FAM | Fig. 4b, Supplementary Fig. 10 |
| Poly ($T_{20}$)-Poly ($C_5$)-FAM | TTTTTTTTTTTTTTTTTTTTCCCCC-FAM | Fig. 4b, Supplementary Fig. 10 |
| Poly ($T_{25}$)-Poly ($C_5$)-FAM | TTTTTTTTTTTTTTTTTTTTTTTTTCCCCC-FAM | Fig. 4b, Supplementary Fig. 10 |
| Poly ($T_{30}$)-Poly ($C_5$)-FAM | TTTTTTTTTTTTTTTTTTTTTTTTTTTTTTCCCCC-FAM | Fig. 4b, Supplementary Fig. 10 |

| | | |
|---|---|---|
| Poly ($T_{10}$)-Poly ($C_2$)-FAM | TTTTTTTTTTCC-FAM | Fig. 4b, Supplementary Fig. 11 |
| Poly ($T_{10}$)-Poly ($C_7$)-FAM | TTTTTTTTTTCCCCCCC-FAM | Fig. 4b, Supplementary Fig. 11 |
| Poly ($T_{10}$)-Poly ($C_{10}$)-FAM | TTTTTTTTTTCCCCCCCCCC-FAM | Fig. 4b, Supplementary Fig. 11 |
| Poly ($T_{10}$)-Poly ($C_{15}$)-FAM | TTTTTTTTTTCCCCCCCCCCCCCCC-FAM | Fig. 4b, Supplementary Fig. 11 |
| Poly ($T_{10}$)-Poly ($C_{20}$)-FAM | TTTTTTTTTTCCCCCCCCCCCCCCCCCCCC-FAM | Fig. 4b, Supplementary Fig. 11 |
| Poly ($T_{10}$)-Poly ($C_{25}$)-FAM | TTTTTTTTTTCCCCCCCCCCCCCCCCCCCCCCCCC-FAM | Fig. 4b, Supplementary Fig. 12 |
| Poly ($T_{10}$)-Poly ($A_2$)-FAM | TTTTTTTTTTAA-FAM | Fig. 4b, Supplementary Fig. 12 |
| Poly ($T_{10}$)-Poly ($A_7$)-FAM | TTTTTTTTTTAAAAAAA-FAM | Fig. 4b, Supplementary Fig. 12 |
| Poly ($T_{10}$)-Poly ($A_{10}$)-FAM | TTTTTTTTTTAAAAAAAAAA-FAM | Fig. 4b, Supplementary Fig. 12 |
| Poly ($T_{10}$)-Poly ($A_{15}$)-FAM | TTTTTTTTTTAAAAAAAAAAAAAAA-FAM | Fig. 4b, Supplementary Fig. 12 |
| Poly ($T_{10}$)-Poly ($A_{20}$)-FAM | TTTTTTTTTTAAAAAAAAAAAAAAAAAAAA-FAM | Fig. 4b, Supplementary Fig. 12 |
| Poly ($T_{10}$)-Poly ($A_{25}$)-FAM | TTTTTTTTTTAAAAAAAAAAAAAAAAAAAAAAAAA-FAM | Fig. 4b, Supplementary Fig. 12 |
| Poly ($A_{10}$)-DNA (1) | AAAAAAAAAATATCTAACCAGAAAGCCACG | Fig. 5a, Supplementary Figs. 13, 17 |
| Poly ($T_{10}$)-DNA (1) | TTTTTTTTTTTATCTAACCAGAAAGCCACG | Fig. 5a, Supplementary Figs. 13, 17 |
| Poly ($C_{10}$)-DNA (1) | CCCCCCCCCCTATCTAACCAGAAAGCCACG | Fig. 5a, Supplementary Figs. 13, 17 |
| Poly ($G_{10}$)-DNA (1) | GGGGGGGGGGTATCTAACCAGAAAGCCACG | Fig. 5a, Supplementary Figs. 13, 17 |
| Poly ($A_{10}$)-DNA (3) | AAAAAAAAAAGTAGCCCGTG ATGCG | Fig. 5a, Supplementary Figs. 13, 17 |
| Poly ($T_{10}$)-DNA (3) | TTTTTTTTTTGTAGCCCGTG ATGCG | Fig. 5a, Supplementary Figs. 13, 17 |
| Poly ($C_{10}$)-DNA (3) | CCCCCCCCCCGTAGCCCGTG ATGCG | Fig. 5a, Supplementary Figs. 13, 17 |
| Poly ($G_{10}$)-DNA (3) | GGGGGGGGGGGTAGCCCGTG ATGCG | Fig. 5a, Supplementary Figs. 13, 17 |
| Padlock probe (with $A_{40}$) | AAGAACTATATTGAAAAAAAAAAAAAAAAAAAAAAAAAAAAAAAAAAAAAAAACACCTTCTCA | Fig. 6a-f |

| | | |
|---|---|---|
| Padlock probe (random) | AAGAACTATATTGAGCTTACAGTGCAGCTG GATCGTAGTGGCAAGCGTTGTCACACCTTC TCA | Fig. 6a-c |
| CFTR G542X Locus (Mutant type) | GACAATATAGTTCTTTGAGAAGGTG | Fig. 6a-f |
| CFTR G542X Locus (Wild type) | GACAATATAGTTCTTGGAGAAGGTG | Fig. 6d-f |
| sgDNA primer-F | GAAATTAATACGACTCACTATAGGGAGATAC GTTGCGTCCGTGATGTTTTAGAGCTAGAAAT AGCAAG | Fig. 6g-o, Supplementary Fig. 16a, b |
| sgDNA primer-R-poly ($A_{17}$) | AAAAAAAAAAAAAAAAAGTAGGTGGCAA GCGTTGTCAGCACCGACTCGGTGCCACTTT TTCAAGTTGATAACGGACTAGCCTTATTTTA ACTTGCTATTTCTAGC | Fig. 6g-o, Supplementary Fig. 16a, b |
| sgDNA primer-R | GTAGGTGGCAAGCGTTGTCAGCACCGACTC GGTGCCACTTTTTCAAGTTGATAACGGACT AGCCTTATTTTAACTTGCTATTTCTAGC | Fig. 6j |
| N gene-primer-F | GAAATTAATACGACTCACTATAGGGATGTCT GATAATGGACCCCA | Fig. 6g-l |
| N gene-primer-R-poly ($A_{15}$) | AAAAAAAAAAAAAAATTAGGCCTGAGTTG AGTCA | Fig. 6g-l |
| N gene-primer-R | TTAGGCCTGAGTTGAGTCA | Fig. 6j |
| Capture DNA (ASFV) | ATCACGGACGCAACGTATCTTTTT-Bio | Fig. 6n, o |
| ASFV-primer-F | Bio-ATGGATACTGAGGGAATAGC | Fig. 6 n, o, Supplementary Fig. 16a, b |
| ASFV-primer-R | CTTACCGATGAAAATGATAC | Fig. 6n, o, Supplementary Fig. 16a, b |
| SH-DNA | SH-CGTAAGTCTGGATCG | Supplementary Fig. 1a, 6a-e, 14b |
| SH-DNA-FAM | SH-TTTTTTTTTT-FAM | Supplementary Fig. 5c, d |
| Capture DNA | Bio-CCTGCTCTAGCAATG | Supplementary Fig. 6a, b |
| Poly ($T_5$)-DNA (hybrid) | TTTTTCATTGCTAGAGCAGG | Supplementary Fig. 6a |
| SH-DNA (hybrid) | SH-CATTGCTAGAGCAGG | Supplementary Fig. 6b |
| Poly ($A_5$)-DNA (5′) | AAAAACGTAAGTCTGGATCG | Supplementary Fig. 7a |
| Poly ($A_{15}$)-DNA (5′) | AAAAAAAAAAAAAAACGTAAGTCTGGATC G | Supplementary Fig. 7a |
| Poly ($A_{20}$)-DNA (5′) | AAAAAAAAAAAAAAAAAAAACGTAAGTCT GGATCG | Supplementary Fig. 7a |
| Poly ($A_{25}$)-DNA (5′) | AAAAAAAAAAAAAAAAAAAAAAAAACGT AAGTCTGGATCG | Supplementary Fig. 7a |
| Poly ($A_{30}$)-DNA (5′) | AAAAAAAAAAAAAAAAAAAAAAAAAAAA AACGTAAGTCTGGATCG | Supplementary Fig. 7a |
| Poly ($A_5$)-DNA (3′) | GCTAGGTCTGAATGCAAAAA | Supplementary Fig. 7a |
| Poly ($A_{15}$)-DNA (3′) | GCTAGGTCTGAATGCAAAAAAAAAAAAAA A | Supplementary Fig. 7a |
| Poly ($A_{20}$)-DNA (3′) | GCTAGGTCTGAATGCAAAAAAAAAAAAAA AAAAAA | Supplementary Fig. 7a |

| | | |
|---|---|---|
| Poly ($A_{25}$)-DNA (3′) | GCTAGGTCTGAATGCAAAAAAAAAAAAAAAAAAAAAAAAA | Supplementary Fig. 7a |
| Poly ($A_{30}$)-DNA (3′) | GCTAGGTCTGAATGCAAAAAAAAAAAAAAAAAAAAAAAAAAAAAA | Supplementary Fig. 7a |
| Poly ($A_5$)-DNA-M | CGCGTTGAATCTATCAAAAACGTAAGTCTGGATCG | Supplementary Fig. 7a |
| Poly ($A_{15}$)-DNA-M | CGCGTTGAATCTATCAAAAAAAAAAAAAAACGTAAGTCTGGATCG | Supplementary Fig. 7a |
| Poly ($A_{20}$)-DNA-M | CGCGTTGAATCTATCAAAAAAAAAAAAAAAAAAAACGTAAGTCTGGATCG | Supplementary Fig. 7a |
| Poly ($A_{25}$)-DNA-M | CGCGTTGAATCTATCAAAAAAAAAAAAAAAAAAAAAAAAACGTAAGTCTGGATCG | Supplementary Fig. 7a |
| Poly ($A_{30}$)-DNA-M | CGCGTTGAATCTATCAAAAAAAAAAAAAAAAAAAAAAAAAAAAAACGTAAGTCTGGATCG | Supplementary Fig. 7a |
| Poly ($C_5$)-DNA (5′) | CCCCCCGTAAGTCTGGATCG | Supplementary Fig. 7b |
| Poly ($C_{15}$)-DN (5′) | CCCCCCCCCCCCCCCCGTAAGTCTGGATCG | Supplementary Fig. 7b |
| Poly ($C_{20}$)-DNA (5′) | CCCCCCCCCCCCCCCCCCCCCGTAAGTCTGGATCG | Supplementary Fig. 7b |
| Poly ($C_{25}$)-DNA (5′) | CCCCCCCCCCCCCCCCCCCCCCCCCCGTAAGTCTGGATCG | Supplementary Fig. 7b |
| Poly ($C_{30}$)-DNA (5′) | CCCCCCCCCCCCCCCCCCCCCCCCCCCCCCCGTAAGTCTGGATCG | Supplementary Fig. 7b |
| Poly ($C_5$)-DNA (3′) | GCTAGGTCTGAATGCCCCCC | Supplementary Fig. 7b |
| Poly ($C_{15}$)-DNA (3′) | GCTAGGTCTGAATGCCCCCCCCCCCCCCCC | Supplementary Fig. 7b |
| Poly ($C_{20}$)-DNA (3′) | GCTAGGTCTGAATGCCCCCCCCCCCCCCCCCCCCC | Supplementary Fig. 7b |
| Poly ($C_{25}$)-DNA (3′) | GCTAGGTCTGAATGCCCCCCCCCCCCCCCCCCCCCCCCCC | Supplementary Fig. 7b |
| Poly ($C_{30}$)-DNA (3′) | GCTAGGTCTGAATGCCCCCCCCCCCCCCCCCCCCCCCCCCCCCCC | Supplementary Fig. 7b |
| Poly ($C_5$)-DNA-M | CGCGTTGAATCTATCCCCCCCGTAAGTCTGGATCG | Supplementary Fig. 7b |
| Poly ($C_{15}$)-DNA-M | CGCGTTGAATCTATCCCCCCCCCCCCCCCCCGTAAGTCTGGATCG | Supplementary Fig. 7b |
| Poly ($C_{20}$)-DNA-M | CGCGTTGAATCTATCCCCCCCCCCCCCCCCCCCCCCGTAAGTCTGGATCG | Supplementary Fig. 7b |
| Poly ($C_{25}$)-DNA-M | CGCGTTGAATCTATCCCCCCCCCCCCCCCCCCCCCCCCCCCGTAAGTCTGGATCG | Supplementary Fig. 7b |
| Poly ($C_{30}$)-DNA-M | CGCGTTGAATCTATCCCCCCCCCCCCCCCCCCCCCCCCCCCCCCCGTAAGTCTGGATCG | Supplementary Fig. 7b |
| Poly ($G_5$)-DNA (5′) | GGGGGCGTAAGTCTGGATCG | Supplementary Fig. 7c |
| Poly ($G_5$)-DNA (3′) | GCTAGGTCTGAATGCGGGGG | Supplementary Fig. 7c |
| Poly ($G_5$)-DNA-M | CGCGTTGAATCTATCGGGGGCGTAAGTCTGGATCG | Supplementary Fig. 7c |
| $T_{10}$-DNA-$T_{10}$ | TTTTTTTTTTCGTAAGTCTGGATCGTTTTTTTTTT | Supplementary Fig. 8a |
| Hairpin DNA1 | TTTTTTTCGCGTTGAATCTATCAAAAAAA | Supplementary Fig. 8b, c |
| Hairpin DNA2 | TTTTTTTCGTAAGTCTGGATCGAAAAAAA | Supplementary Fig. 8c |
| Hairpin DNA3 | TTTTTTTGTAGCCCGTGATGCGAAAAAAA | Supplementary Fig. 8c |

| | | |
|---|---|---|
| Hairpin DNA4 | TTTTTTTGATCCACCTGCACTGTAAGCAAA AAAA | Supplementary Fig. 8c |
| Hairpin DNA5 | TTTTTTTATCTAACCAGAAAGCCACGAAA AAAA | Supplementary Fig. 8c |
| Hairpin DNA6 | TTTTTTTGACAACGCTTGCCACCTACAAAA AAA | Supplementary Fig. 8c |
| Poly ($A_{10}$)-DNA (4) | AAAAAAAAAAGATCCACCTGCACTGTAAG C | Supplementary Figs. 13, 17 |
| Poly ($T_{10}$)-DNA (4) | TTTTTTTTTTGATCCACCTGCACTGTAAGC | Supplementary Figs. 13, 17 |
| Poly ($C_{10}$)-DNA (4) | CCCCCCCCCCGATCCACCTGCACTGTAAGC | Supplementary Figs. 13, 17 |
| Poly ($G_{10}$)-DNA (4) | GGGGGGGGGGGATCCACCTGCACTGTAAG C | Supplementary Figs. 13, 17 |
| Poly ($A_{10}$)-DNA (5) | AAAAAAAAAACGCGTTGAATCTATC | Supplementary Figs. 13, 17 |
| Poly ($T_{10}$)-DNA (5) | TTTTTTTTTTCGCGTTGAATCTATCA | Supplementary Figs. 13, 17 |
| Poly ($C_{10}$)-DNA (5) | CCCCCCCCCCCGCGTTGAATCTATC | Supplementary Figs. 13, 17 |
| Poly ($G_{10}$)-DNA (5) | GGGGGGGGGGCGCGTTGAATCTATC | Supplementary Figs. 13, 17 |
| Poly ($A_{10}$)-DNA (6) | AAAAAAAAAAGACAACGCTTGCCACCTAC | Supplementary Figs. 13, 17 |
| Poly ($T_{10}$)-DNA (6) | TTTTTTTTTTGACAACGCTTGCCACCTAC | Supplementary Figs. 13, 17 |
| Poly ($C_{10}$)-DNA (6) | CCCCCCCCCCGACAACGCTTGCCACCTAC | Supplementary Figs. 13, 17 |
| Poly ($G_{10}$)-DNA (6) | GGGGGGGGGGGACAACGCTTGCCACCTAC | Supplementary Figs. 13, 17 |
| Poly ($T^*_5$)-DNA | T*T*T*T*T*CATTGCTAGAGCAGG | Supplementary Fig. 15b |
| Poly ($T^*_7$)-DNA | T*T*T*T*T*T*T*CATTGCTAGAGCAGG | Supplementary Fig. 15b |
| Poly ($T^*_{10}$)-DNA | T*T*T*T*T*T*T*T*T*T*CATTGCTAGAGCAG G | Supplementary Fig. 15b |
| Poly ($T^*_{15}$)-DNA | T*T*T*T*T*T*T*T*T*T*T*T*T*T*T*CATTGC TAGAGCAGG | Supplementary Fig. 15b |
| Poly ($T^*_7$)-DNA-Poly ($T_7$) | T*T*T*T*T*T*T*CATTGCTAGAGCAGGTTTT TTT | Supplementary Fig. 15b |

[a]M stands for the middle region. The mutant base is highlighted in red. * stands for the phosphorothioate (PS) modification.

**Supplementary Table 3** Genomic sequences used in this work.

| Names | Sequences (from 5′ to 3′) |
|---|---|
| N gene of SARS-CoV-2 | ATGTCTGATAATGGACCCCAAAATCAGCGAAATGCACCCCGCA<br>TTACGTTTGGTGGACCCTCAGATTCAACTGGCAGTAACCAGAA<br>TGGAGAACGCAGTGGGGCGCGATCAAAACAACGTCGGCCCCA<br>AGGTTTACCCAATAATACTGCGTCTTGGTTCACCGCTCTCACTC<br>AACATGGCAAGGAAGACCTTAAATTCCCTCGAGGACAAGGCG<br>TTCCAATTAACACCAATAGCAGTCCAGATGACCAAATTGGCTAC<br>TACCGAAGAGCTACCAGACGAATTCGTGGTGGTGACGGTAAA<br>ATGAAAGATCTCAGTCCAAGATGGTATTTCTACTACCTAGGAAC<br>TGGGCCAGAAGCTGGACTTCCCTATGGTGCTAACAAAGACGGC<br>ATCATATGGGTTGCAACTGAGGGAGCCTTGAATACACCAAAAG<br>ATCACATTGGCACCCGCAATCCTGCTAACAATGCTGCAATCGTG<br>CTACAACTTCCTCAAGGAACAACATTGCCAAAAGGCTTCTACG<br>CAGAAGGGAGCAGAGGCGGCAGTCAAGCCTCTTCTCGTTCCT<br>CATCACGTAGTCGCAACAGTTCAAGAAATTCAACTCCAGGCAG<br>CAGTAGGGGAACTTCTCCTGCTAGAATGGCTGGCAATGGCGGT<br>GATGCTGCTCTTGCTTTGCTGCTGCTTGACAGATTGAACCAGCT<br>TGAGAGCAAAATGTCTGGTAAAGGCCAACAACAACAAGGCCA<br>AACTGTCACTAAGAAATCTGCTGCTGAGGCTTCTAAGAAGCCT<br>CGGCAAAAACGTACTGCCACTAAAGCATACAATGTAACACAAG<br>CTTTCGGCAGACGTGGTCCAGAACAAACCCAAGGAAATTTTG<br>GGGACCAGGAACTAATCAGACAAGGAACTGATTACAAACATT<br>GGCCGCAAATTGCACAATTTGCCCCCAGCGCTTCAGCGTTCTT<br>CGGAATGTCGCGCATTGGCATGGAAGTCACACCTTCGGGAACG<br>TGGTTGACCTACACAGGTGCCATCAAATTGGATGACAAAGATC<br>CAAATTTCAAAGATCAAGTCATTTTGCTGAATAAGCATATTGAC<br>GCATACAAAACATTCCCACCAACAGAGCCTAAAAAGGACAAA<br>AAGAAGAAGGCTGATGAAACTCAAGCCTTACCGCAGAGACAG<br>AAGAAACAGCAAACTGTGACTCTTCTTCCTGCTGCAGATTTGG<br>ATGATTTCTCCAAACAATTGCAACAATCCATGAGCAGTGCTGA<br>CTCAACTCAGGCCTAA |
| VP72 gene fragment of ASFV | AAGGTTCACGTTCTCGTTAAACCAAAAGCGCAGCTTAATCCAG<br>AGCGCAAGAGGGGGCTGATAGTATTTAGGGGTTTGAGGTCCAT<br>TACAGCTGTAATGAACATTACGTCTTATGTCCAGATACGTTGCG<br>TCCGTGATAGGAGTGATATCTTGTTTACCTGCTGTTTGGATATTG<br>TGAGAGTTCTCGGGAAAATGTTGTGAAAGAAATTTCGGGTTGG<br>TATGGCTGCACGTTCGCTGCGTATCATTTTCATCGGTAAG |
